# KDM4 inhibition alters H3K9me3 distribution and developmental responses in SETDB1-deficient embryonic stem cells

**DOI:** 10.64898/2026.10.14.758827

**Authors:** Rui Geng, Benjamin L. Kidder

## Abstract

SETDB1 contributes to histone H3 lysine 9 methylation, whereas KDM4 demethylases act on H3K9 and, for KDM4A-C, H3K36. We examined whether Setdb1 status modifies responses to the KDM4-family inhibitor QC6352 in embryonic stem cells (ESCs) and embryoid bodies (EBs). QC6352 shifted Setdb1-depleted ESC colonies toward more compact morphology and increased alkaline-phosphatase (AP) staining, but also affected Setdb1-intact cells. Separate Kdm4a, Kdm4b, and Kdm4c knockdowns identified overlapping and member-specific transcriptional responses. Factorial RNA-seq revealed broadly concordant transcriptional responses to KDM4 inhibition in Setdb1-intact and depleted ESCs, alongside a smaller subset of genes with genotype-dependent treatment effects. Within this subset, two gene modules showed attenuation or reversal of Setdb1-associated expression changes following KDM4 inhibition. ChIP-seq revealed changes in the relative distribution of H3K9me3, H3K36me3, and H3K4me3 between promoters and gene bodies following KDM4 inhibition. KDM4 inhibition altered H3K9me3 distributions across repeat families in both Setdb1-intact and depleted ESCs. QC6352 also altered EB morphology and produced transcriptional responses that differed between ESCs and EBs. These findings show that KDM4 inhibition remodels chromatin and transcription in both Setdb1-intact and depleted ESCs, with Setdb1 status influencing a subset of transcriptional responses.

## INTRODUCTION

Embryonic stem cells (ESCs) sustain self-renewal while retaining the capacity to enter diverse developmental lineages^1–4^. This plasticity depends on chromatin systems that preserve genome integrity without preventing rapid transcriptional reconfiguration, including covalent histone modifications^5–8^. Histone H3 lysine 9 methylation (H3K9me) is a central component of this regulatory architecture^9–13^. Its trimethylated form, H3K9me3, is enriched within heterochromatin and contributes to repression of endogenous retroviruses (ERVs)^14, 15^. Yet H3K9me3 is not static: redistribution of the mark accompanies ESC-state transitions and can alter the accessibility of lineage-associated programs^16, 17^.

H3K9 methylation is controlled by opposing enzymatic activities. Setdb1 (ESET/KMT1E) contributes to H3K9 mono-, di-, and trimethylation and is required for heterochromatin maintenance and ERV restriction in ESCs^5, 15, 18^. Demethylation provides an opposing mechanism through which these repressive states can be remodeled^19, 20^. KDM4A-C remove di- and trimethyl groups from H3K9 and H3K36, whereas KDM4D acts on H3K9 without the corresponding H3K36 activity; these substrate differences constrain interpretation of family-directed perturbations^19–24^. Individual family members can also influence enhancer and lineage programs^25^. Their partially overlapping substrate specificities and functions, however, make it difficult to infer whether a phenotype produced by a family-directed inhibitor reflects one paralog, combined activity of several paralogs, or broader pharmacologic effects^26^. SETDB1 acts within a broader network of H3K9 methyltransferases that includes SUV39H1, SUV39H2, EHMT1/GLP, EHMT2/G9a, and SETDB2, each with distinct methylation-state and genomic preferences. This network provides a basis for H3K9 methylation to persist following Setdb1 depletion, allowing KDM4 inhibition to influence the turnover and distribution of residual methylated chromatin.

How Setdb1-dependent methylation and KDM4-dependent demethylation interact to stabilize ESC identity therefore remains unresolved^27^. Perturbing this axis can influence enhancers, developmental transcription, and higher-order chromatin organization^28^, but existing loss-of-function studies also indicate that KDM4-family responses are distributed across partially redundant enzymes^26^. Small-molecule KDM4 inhibitors, including QC6352, provide a complementary means to suppress family activity^29–32^. Most studies of these compounds have focused on cancer models^31, 32^, leaving several unanswered questions especially important in pluripotent cells: whether KDM4 inhibition can buffer Setdb1 deficiency; whether a large inhibitor-induced transcriptome is reproducible across Setdb1 genotypes; whether KDM4 inhibition alters the abundance and genomic organization of histone modifications; and whether the resulting response persists or changes during differentiation.

We addressed these questions using a balanced genotype-by-treatment design that combined conditional Setdb1 depletion with KDM4 inhibition in ESCs and differentiating embryoid bodies. Paralog-resolved shRNA experiments defined overlapping and member-specific transcriptional consequences of KDM4A, KDM4B, and KDM4C depletion, while factorial RNA-seq separated conserved QC6352 responses from formal Setdb1-dependent interactions. Multi-mark ChIP-seq quantified promoter-to-gene-body redistribution, and repeat-family analyses evaluated whether H3K9me3 remodeling predicted repeat transcription. The results reveal a large but highly reproducible KDM4i response that is mostly conserved across Setdb1 backgrounds, accompanied by changes in selected KO cellular features, relative histone-mark distributions, and developmental-stage-dependent transcriptional responses. These findings support a model in which perturbation of KDM4 activity reorganizes the distribution and developmental interpretation of methylated chromatin rather than simply reversing Setdb1-dependent chromatin states.

## RESULTS

### KDM4 inhibition alters colony morphology and AP staining in Setdb1-depleted ESCs

Setdb1 and KDM4 enzymes have opposing activities in H3K9 methylation turnover, raising the possibility that inhibiting KDM4-mediated demethylation could buffer the loss of Setdb1-dependent methylation. We tested this possibility using a 2 × 2 factorial design comparing Setdb1 conditional-control (cKO) and Setdb1-depleted (KO) ESCs treated with DMSO or the KDM4 inhibitor QC6352 (KDM4i; **Fig. 1A**). Setdb1 loss disrupted compact ESC colony architecture, whereas KDM4i shifted KO cultures toward more compact colony features and produced a concentration-dependent morphological response in R1 ESCs and Setdb1 cKO/KO cells (**Fig. 1B; Supplemental Fig. S1A-B**). SETDB1 depletion was confirmed by immunoblotting (**Fig. 1C**). These findings indicate that the Setdb1-deficient ESC phenotype remained partially responsive to KDM4 inhibition.

**Figure 1.**
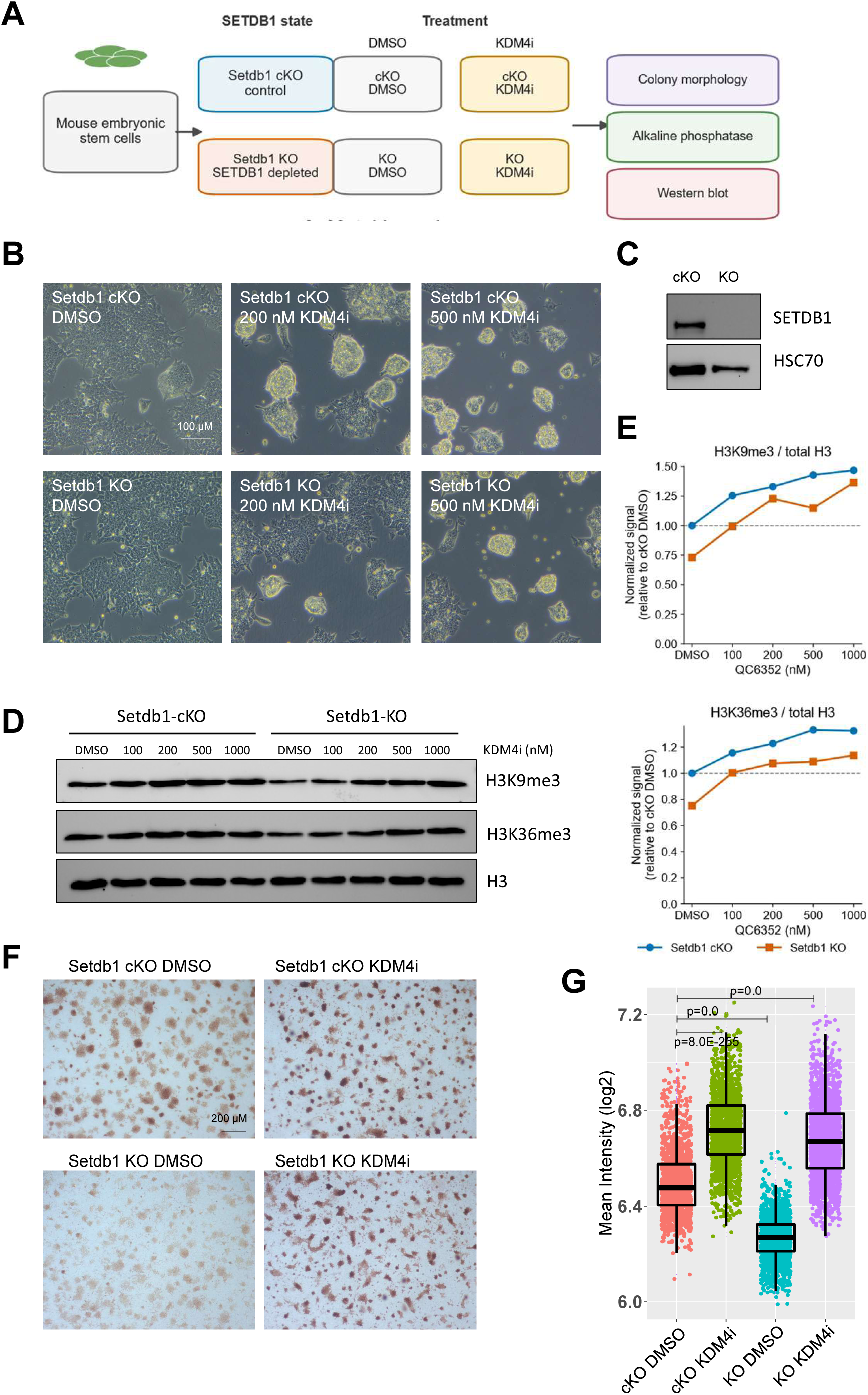
KDM4 inhibition alters colony morphology AP staining and histone methylation in Setdb1-depleted ESCs. **(A)** Experimental design comparing Setdb1 cKO and KO ESCs treated with DMSO or KDM4i and assessed by colony morphology, AP staining, and immunoblotting. **(B)** Representative bright-field images at the indicated KDM4i concentrations. **(C)** SETDB1 immunoblot; HSC70 is the loading control. **(D)** H3K9me3, H3K36me3, and total-H3 immunoblots across the KDM4i dose series. **(E)** Densitometric H3K9me3/total-H3 and H3K36me3/total-H3 ratios normalized to Setdb1 cKO DMSO. **(F)** Representative AP-stained cultures. **(G)** Object-level log2 mean AP intensity. Boxes denote the interquartile range and the center line denotes the median. Experiments were performed with at least three to four independent biological replicates. Each point in G represents an individual colony sampled from the replicate cultures.

H3K9me3 and H3K36me3 immunoblot signals increased relative to total H3 across the KDM4i dose series in both genotypes, although baseline levels and response magnitude differed between cKO and KO cells (**Fig. 1D-E**).

An alkaline-phosphatase (AP) assay assessed a cell-state-associated marker. Setdb1 KO cultures exhibited reduced AP staining under DMSO, whereas QC6352 increased AP-positive colony staining (**Fig. 1F-G**). Dose-response imaging and object-level quantification in R1 ESCs showed a similar trend (**Supplemental Fig. S1C-D**). Thus, QC6352 shifted selected features of KO cultures toward the control phenotype, while also affecting Setdb1-intact cells.

### KDM4-family perturbations converge on a shared program while retaining member-specific functions

We next asked whether depletion of individual KDM4 paralogs produced overlapping or distinct transcriptional responses that could provide genetic context for the pharmacologic phenotype. We therefore used independent shRNAs targeting Kdm4a, Kdm4b, or Kdm4c. qRT-PCR confirmed target reduction relative to shLuc across multiple independent shRNAs targeting Kdm4a, Kdm4b, or Kdm4c (**Fig. 2A**). Each shRNA construct was independently profiled by RNA-seq, allowing target-associated responses to be evaluated across multiple hairpins. Immunoblotting independently confirmed depletion of KDM4A, KDM4B, and KDM4C across the tested shRNA constructs, with variable knockdown efficiency among individual hairpins (**Supplemental Fig. S2A**).

**Figure 2.**
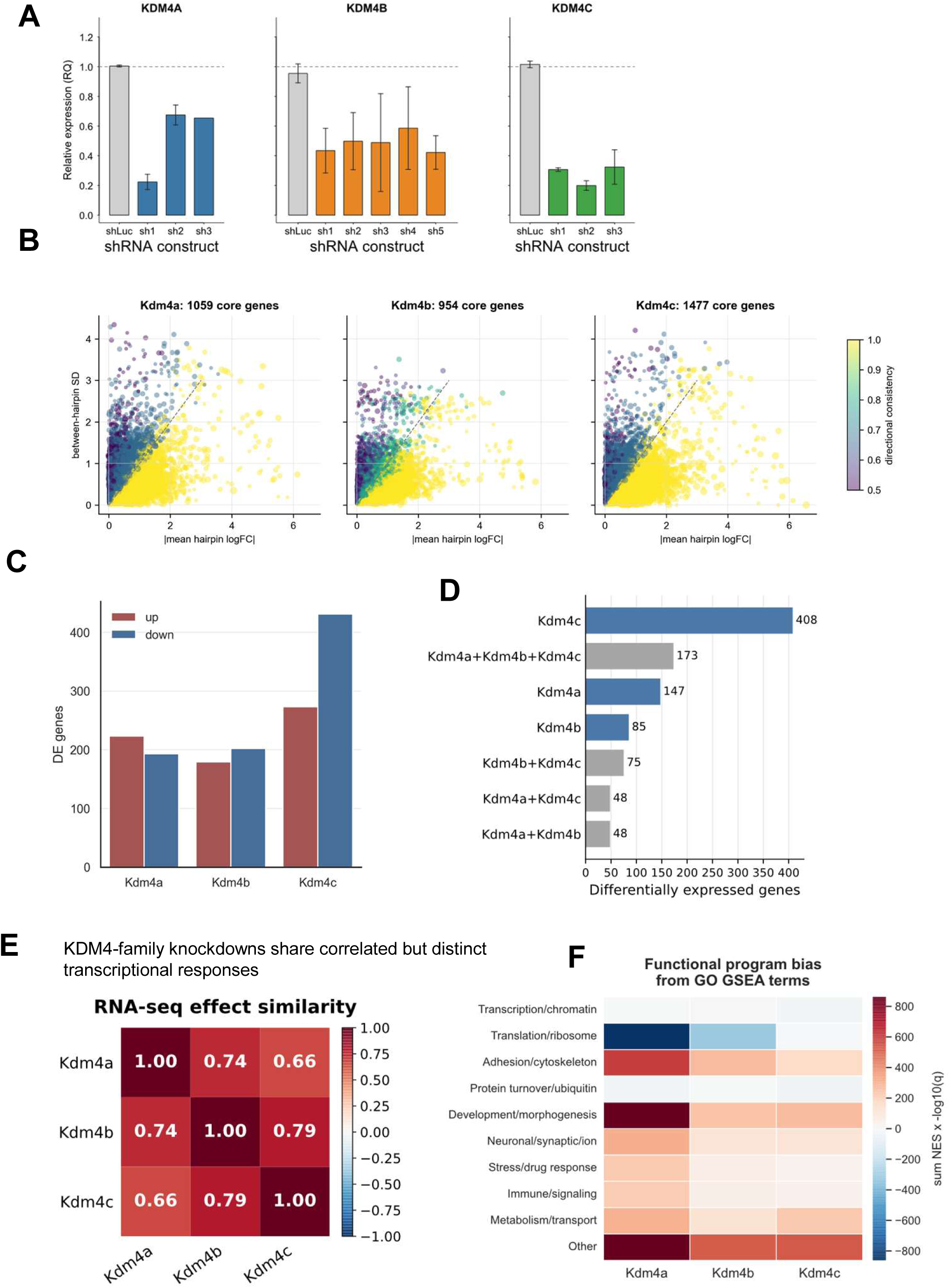
Independent KDM4-family knockdowns reveal overlapping and member-specific ESC transcriptional programs. **(A)** qRT-PCR expression of Kdm4a, Kdm4b, and Kdm4c after the indicated shRNAs, relative to shLuc. **(B)** Gene-level mean absolute hairpin effect versus between-hairpin variability. Density shows all genes; highlighted points meet the core-consensus definition (|mean log2FC| ≥ 0.5, complete directional agreement, and significance in at least half of target-specific hairpins). **(C)** Up- and downregulated gene counts for each aggregate family-member knockdown response derived from independently analyzed shRNAs. **(D)** Shared and member-specific differentially expressed gene sets. **(E)** Pairwise correlation of genome-wide RNA-seq effects among Kdm4a, Kdm4b, and Kdm4c knockdowns. **(F)** Representative GO GSEA programs. Color encodes signed NES, point size encodes −log10(BH-FDR), and dark outlines denote BH-FDR < 0.05.

Integrating RNA-seq effects across the independently analyzed shRNAs identified 1,059 Kdm4a, 954 Kdm4b, and 1,477 Kdm4c core-consensus genes (Fig. 2B). Kdm4c knockdown produced the largest differential-expression burden (**Fig. 2C**), yet the response was not simply paralog-specific: 173 genes were shared by all three knockdowns, whereas 147, 85, and 408 genes were unique to Kdm4a, Kdm4b, and Kdm4c, respectively (**Fig. 2D**). Genome-wide effects were positively correlated, most strongly between Kdm4b and Kdm4c (r = 0.79), followed by Kdm4a-Kdm4b (r = 0.74) and Kdm4a-Kdm4c (r = 0.66; **Fig. 2E**). These data identify overlapping KDM4A-, KDM4B-, and KDM4C-responsive transcriptional programs superimposed on substantial member-specific effects.

Functional enrichment reinforced this division of labor. Translation/ribosome and RNA-processing programs were negatively enriched, whereas adhesion/cytoskeletal and developmental or morphogenetic programs were positively enriched, with paralog-dependent magnitudes (**Fig. 2F**). PROGENy analysis identified the strongest inferred Hypoxia and EGFR activity in the KDM4A knockdown response (**Supplemental Fig. S2B-C**). The overlapping transcriptional effects of Kdm4a, Kdm4b, and Kdm4c knockdown establish partially shared functions among the tested paralogs, while their distinct effects show that no single knockdown reproduces the broader QC6352 response.

### KDM4 inhibition induces a broad transcriptional response that is largely conserved across Setdb1 genotypes

We next assessed whether the KDM4i transcriptional response was reproducible across Setdb1 backgrounds. Replicates clustered closely, and the four genotype-treatment groups separated clearly by principal-component analysis (**Fig. 3A**), indicating consistent transcriptional responses across the experimental conditions.

**Figure 3.**
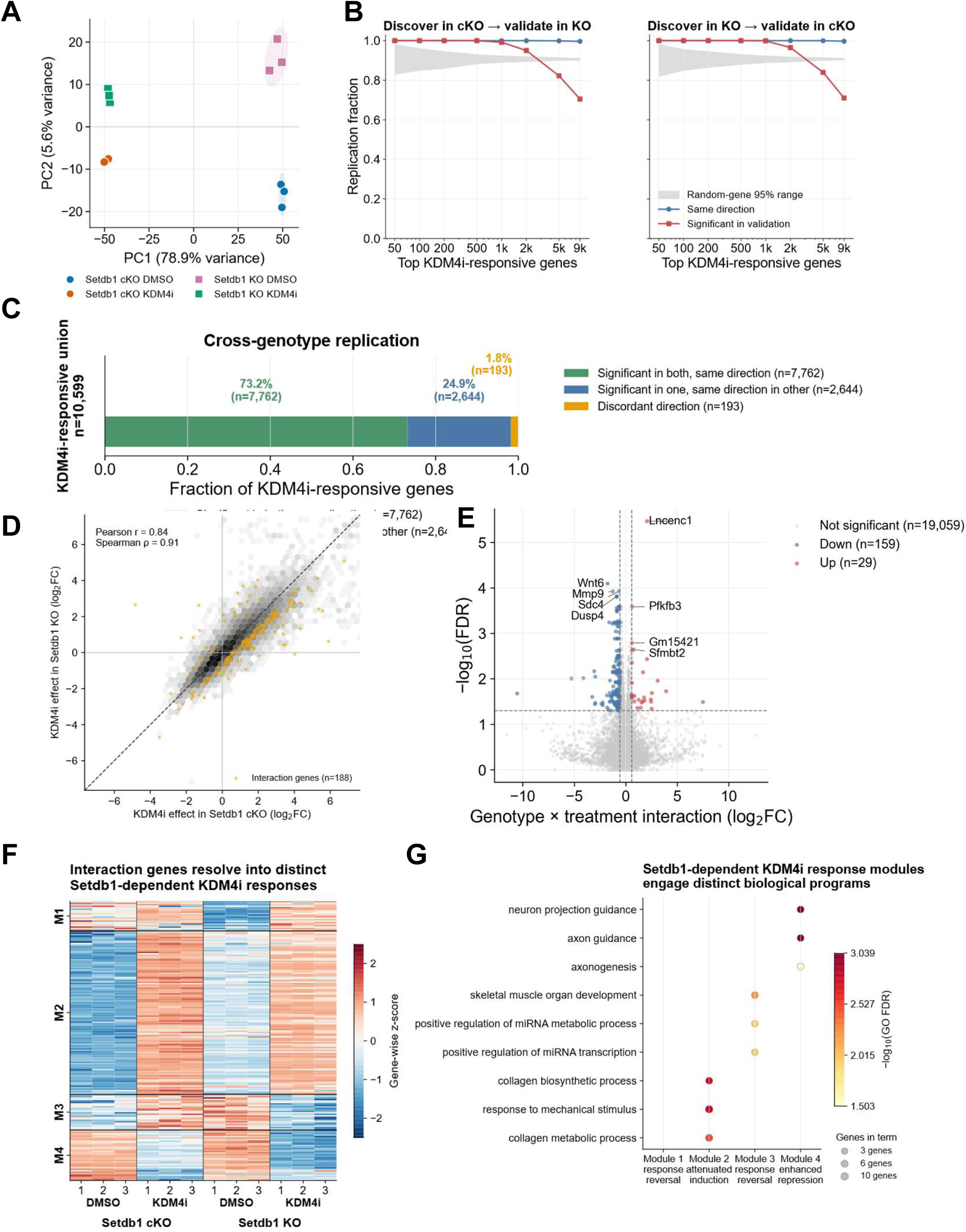
KDM4 inhibition drives a conserved transcriptional program across Setdb1 states. (A) PCA of the 500 most variable expressed genes from three biological replicates per condition; ellipses are descriptive covariance regions. (B) Reciprocal discovery/validation curves. Blue, same direction; red, independently significant in the validation genotype (BH-FDR < 0.05 and |log2FC| ≥ 0.58); gray, 2.5th-97.5th percentile from 300 size-matched random-gene resamples. (C) Classification of the 10,599 genes responsive to KDM4 inhibition in at least one Setdb1 genotype. Of these, 7,762 genes were significant and changed in the same direction in both genotypes, 2,644 were significant in one genotype and changed in the same direction in the other, and 193 changed in opposite directions. Counts and percentages are shown above the bar. (D) KDM4i log2FC in cKO versus KO cells for all 19,247 tested genes; orange points are all 188 formal interaction genes. (E) Interaction volcano plot; significant interaction genes were defined by BH-FDR < 0.05 and |interaction log2FC| ≥ 0.58. (F) Gene-wise z-scored logCPM heatmap of the interaction genes. K-means clustering of gene-wise z-scored profiles (k = 4, 50 starts, seed 20260724) defined four modules: response reversal (M1, n = 20), attenuated induction (M2, n = 110), response reversal (M3, n = 24), and enhanced repression (M4, n = 34). (G) GO Biological Process enrichment of modules using all 19,247 expressed genes as background. Color denotes −log10(BH-FDR) and size denotes module genes in the term.

Reciprocal discovery-validation analysis provided a more stringent test. Genes ranked by the KDM4i response in one genotype were evaluated with an independently fitted contrast in the other genotype (**Fig. 3B**). The top 50-500 genes were 100% directionally concordant and independently significant; even among the top 9,000 genes, 99.6-99.7% retained direction and 70.5-71.0% independently met the validation threshold. These values exceeded the 2.5th-97.5th percentile range from 300 size-matched random-gene resamples. The scale of the inhibitor response is therefore supported by reciprocal replication, not merely by small P values in a single comparison.

At BH-FDR < 0.05 and |log2 fold change| ≥ 0.58, the KDM4i-responsive union contained 10,599 genes (**Fig. 3C**). Of these, 7,762 (73.2%) were significant and concordant in both genotypes, another 2,644 (24.9%) were significant in one genotype and concordant in the other, and only 193 (1.8%) changed in opposing directions. Accordingly, 98.2% of the union retained response polarity across Setdb1 backgrounds. A formal genotype-by-treatment test identified only 188 interaction genes among 19,247 tested genes (29 positive and 159 negative coefficients; **Fig. 3D-E**). Of these, 171 belonged to the KDM4i-responsive union and comprise the 1.6% interaction fraction in **Fig. 3C**; the other 17 did not meet the per-genotype response definition. This resolves the 171-versus-188 counts and, more importantly, shows that Setdb1 status modulates only a restricted edge of an otherwise genotype-conserved KDM4i program.

Effect sizes were correlated between genotypes (Pearson r = 0.84; Spearman ρ = 0.91; **Fig. 3D**). Within the 188 interaction genes, seeded k-means clustering of gene-wise z-scored logCPM profiles across all 12 libraries resolved response-reversal modules M1 (20 genes) and M3 (24 genes), an attenuated-induction module M2 (110 genes), and an enhanced-repression module M4 (34 genes; **Fig. 3F**). The 44 genes in M1 and M3 are candidate correlates of attenuation or reversal of Setdb1-associated expression changes. GO analysis using the expressed-gene background linked the interaction modules to collagen/mechanical-response, miRNA-regulatory, and axon-guidance processes (**Fig. 3G**). The broad conserved response and the restricted genotype-sensitive modules address distinct aspects of the experiment.

### KDM4 inhibition reorganizes the intragenic distribution of multiple histone marks

We next asked whether the conserved transcriptional response was accompanied by a conserved spatial reorganization of chromatin. Input-normalized, expression-stratified metagene profiles revealed mark-specific responses in both Setdb1 backgrounds (**Fig. 4A**). H3K4me3 KDM4i/DMSO ratios were greatest across the bodies of low-expression genes, whereas H3K9me3 accumulation increased with expression and was most pronounced across highly expressed genes. Setdb1 loss alone produced comparatively modest mean changes under DMSO. ChIP-seq quality-control analyses showed mark-specific differences in enriched-region counts (**Supplemental Fig. S3A**), while the genomic distribution of enriched regions remained broadly consistent across conditions (**Supplemental Fig. S3B**). Signal PCA separated genotype-treatment groups for each histone mark (**Supplemental Fig. S3C**), and fixed-bin correlations showed strong within-mark similarity across biological replicates (**Supplemental Fig. S3D**), supporting the reproducibility of the chromatin profiles used for downstream analyses.

**Figure 4.**
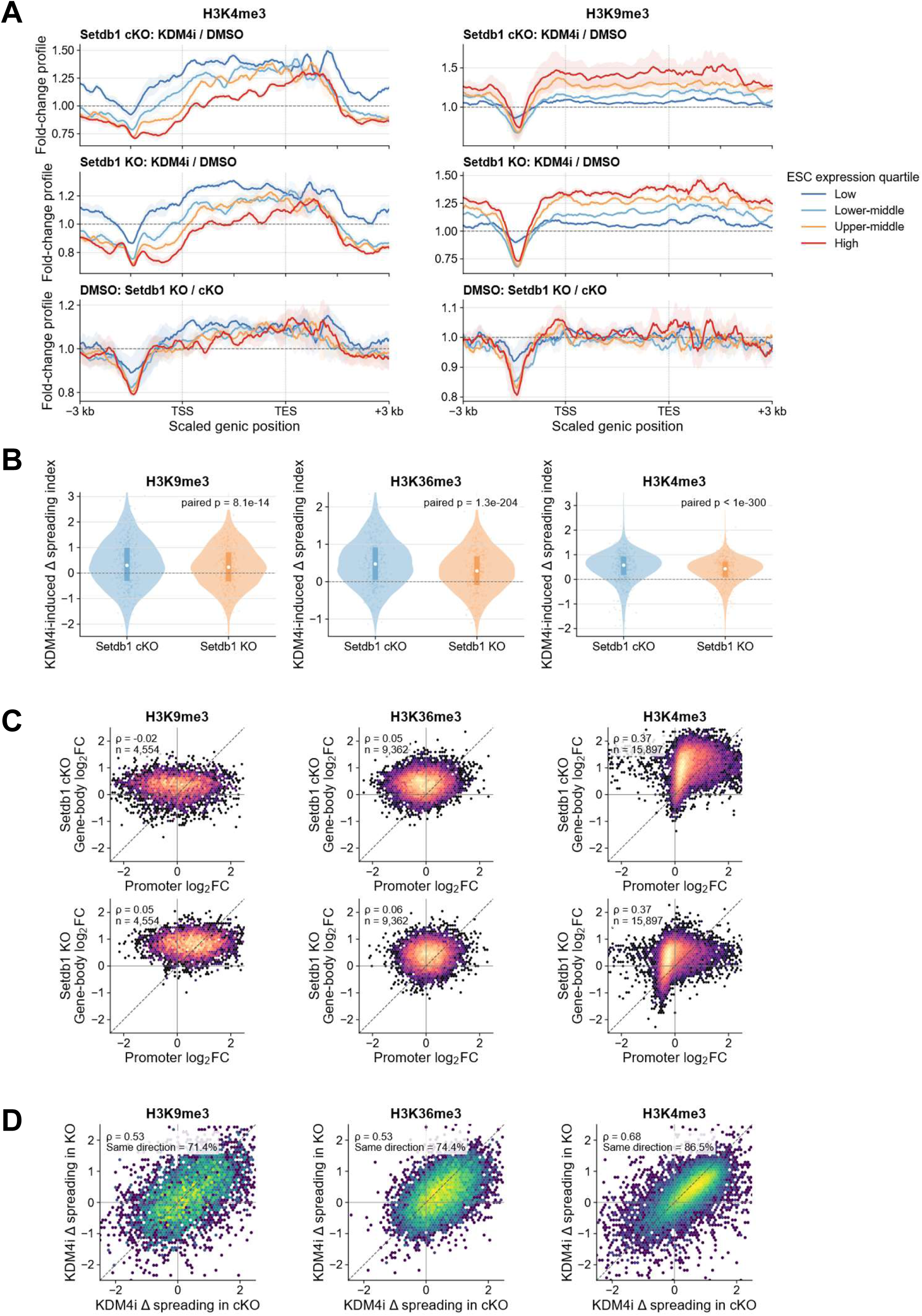
KDM4 inhibition induces expression-dependent and genotype-conserved redistribution of genic chromatin. **(A)** Input-normalized expression-quartile metagene profiles for H3K4me3 and H3K9me3. Lines show replicate means; ribbons span replicate values. Dashed horizontal lines denote ratio 1 and dotted vertical lines mark TSS and TES. **(B)** KDM4i-induced change in gene-body-to-promoter index, defined as Δ log2(gene-body signal/promoter signal). Violin width shows gene density, thick segments show interquartile ranges, and open circles show medians. Displayed paired Wilcoxon tests are descriptive matched-gene tests, not biological-replicate tests (H3K9me3 n = 4,554; H3K36me3 n = 9,362; H3K4me3 n = 15,897). **(C)** Input-normalized promoter log2(KDM4i/DMSO) versus gene-body log2(KDM4i/DMSO) phase portraits in cKO (top) and KO (bottom) cells. The dashed diagonal denotes equal promoter and gene-body response. **(D)** Cross-genotype concordance of KDM4i-induced index changes. Axes use the same Δ-index definition as B; dashed lines denote identity. Spearman correlations and same-direction fractions are shown.

To summarize relative positional signal, we calculated log2(gene-body signal/promoter signal), termed the gene-body-to-promoter index, and its within-genotype QC6352-minus-DMSO change. Median changes were 0.301, 0.473, and 0.575 in cKO cells for H3K9me3, H3K36me3, and H3K4me3, respectively, compared with 0.230, 0.287, and 0.427 in KO cells (**Fig. 4B**). The larger cKO responses do not support preferential chromatin restoration in KO cells. An increased ratio can reflect a gene-body increase, a promoter decrease, or both, and does not establish physical spreading. Paired Wilcoxon tests over matched genes are descriptive rather than inference from independent biological replicates.

Promoter-versus-gene-body phase portraits resolved the ratio into its component signals and placed most genes above the identity line, indicating a larger relative KDM4i effect in gene bodies than promoters (**Fig. 4C**). This response reproduced across genotypes: index changes correlated at ρ = 0.53 for H3K9me3, 0.53 for H3K36me3, and 0.68 for H3K4me3, with same-direction fractions of 71.4%, 74.4%, and 86.5% (**Fig. 4D**). An increase in the H3K36me3 gene-body/promoter ratio is compatible with a decline in total H3K36me3 signal because abundance and intragenic distribution are distinct quantities; relative preservation of gene-body signal or a proportionally larger promoter decrease raises the ratio. A ChIP-only sensitivity analysis that omitted input-DNA normalization showed similar positional patterns for H3K36me3 and H4K20me3. Expression-stratified metagene profiles showed condition-dependent redistribution across promoters and gene bodies (**Supplemental Fig. S4A,C**), while gene-body-to-promoter-index distributions confirmed increased gene-body relative to promoter signal after KDM4 inhibition in both Setdb1 backgrounds (**Supplemental Fig. S4B,D**). Together, the input-normalized and ChIP-only analyses identify reproducible, mark-specific positional chromatin reorganization as a consequence of KDM4 inhibition.

### Repeat-associated H3K9me3 remodeling is reproducible but only partially aligned with repeat transcription

Because Setdb1-dependent heterochromatin contributes to repeat regulation, we integrated H3K9me3 density with repeat-associated RNA across 49 repeat families represented in both biological replicates and all four ESC conditions. KDM4i broadly altered repeat-associated H3K9me3, with heterogeneous family-level responses that were generally larger in cKO than KO cells (**Fig. 5A**). These effects were highly reproducible across biological replicates in both genotypes (Spearman ρ = 0.84; **Fig. 5B**), indicating that the ranked family responses were not driven by isolated measurements.

**Figure 5.**
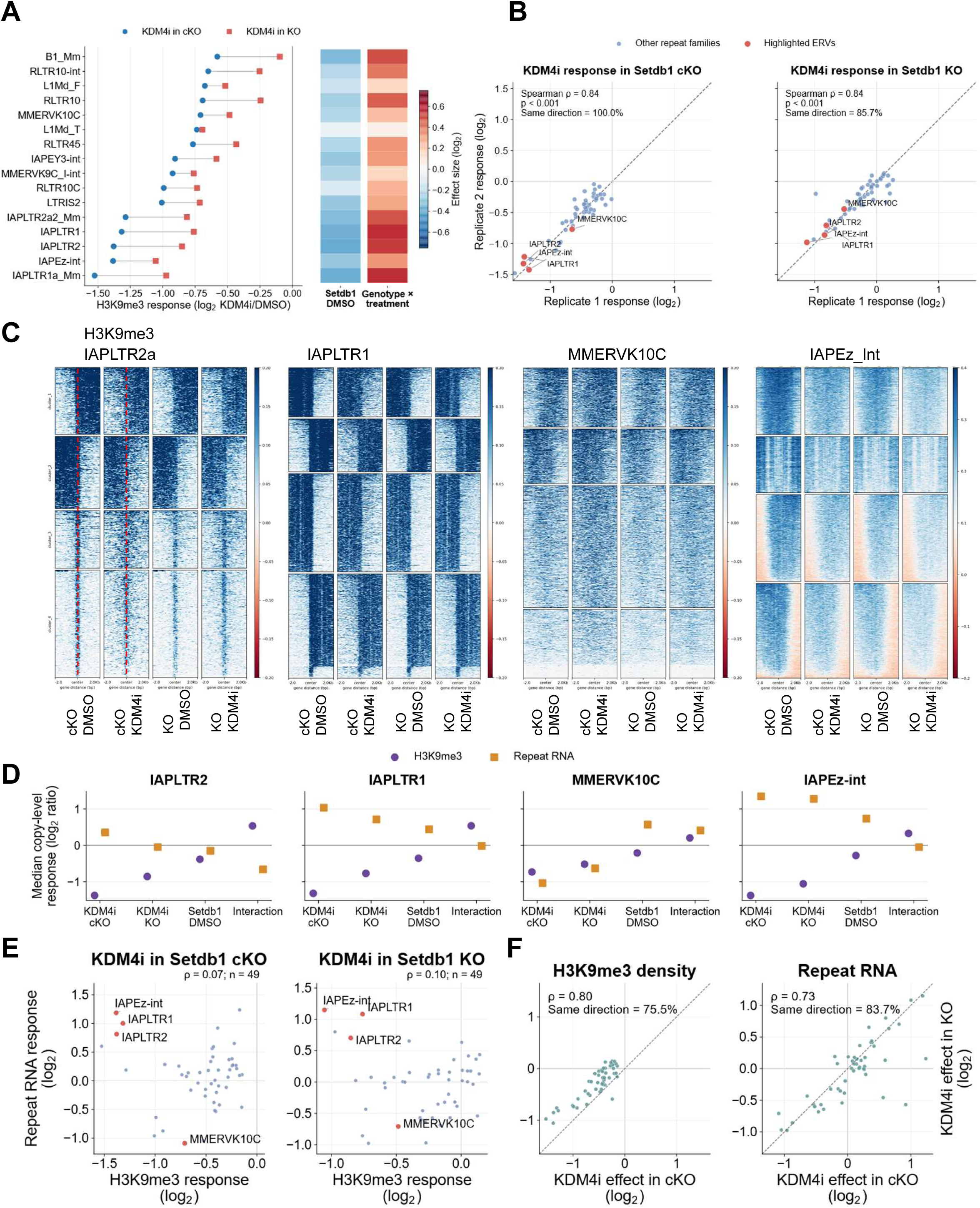
KDM4 inhibition reproducibly remodels repeat-associated H3K9me3 with limited coupling to repeat transcription. **(A)** Repeat-family median H3K9me3 responses to KDM4i in cKO and KO cells; adjacent tiles show the Setdb1 effect under DMSO and the genotype × treatment interaction. **(B)** Biological-replicate concordance across 49 repeat families. Red points denote highlighted ERV families and dashed lines denote identity. **(C)** Element-centered H3K9me3 heatmaps for IAPLTR2a, IAPLTR1, MMERVK10C, and IAPEz-int using one shared color scale. **(D)** Repeat-family median H3K9me3-density effects and family-aggregate repeat-RNA effects for highlighted ERVs. **(E)** Association between KDM4i-induced H3K9me3 and repeat-RNA responses in cKO and KO cells. **(F)** Cross-genotype concordance of KDM4i effects on H3K9me3 density and repeat RNA. Spearman correlations and same-direction fractions are shown.

Element-centered heatmaps further revealed distinct spatial H3K9me3 responses at IAPLTR2a, IAPLTR1, MMERVK10C, and IAPEz-int elements (**Fig. 5C**), with additional repeat classes shown in **Supplemental Fig. S5**. The shared color scale allows direct comparison across the four highlighted families. **Figure 5D** summarizes, for each selected family, the median H3K9me3 response together with the corresponding family-level RNA response.

At selected ERV families, including IAPLTR2, IAPLTR1, and IAPEz-int, reduced H3K9me3 was accompanied by increased repeat RNA, consistent with the repressive role of H3K9me3. However, this inverse relationship was not consistent across all 49 repeat families. Across families, H3K9me3 and RNA responses showed only weak overall association within each genotype (ρ = 0.07 in cKO and ρ = 0.10 in KO; **Fig. 5E**). By contrast, each response type was conserved across Setdb1 backgrounds, with cross-genotype correlations of ρ = 0.80 for H3K9me3 and ρ = 0.73 for repeat RNA and same-direction agreement of 75.5% and 83.7%, respectively (**Fig. 5F**).

Selected ERV families showed reduced relative H3K9me3 signal together with increased repeat RNA, but the weak association across the 49 analyzed families indicates that these signal changes are not a reliable family-wide predictor of RNA response. The larger cKO chromatin responses do not support preferential restoration in KO cells.

### KDM4 inhibition alters embryoid-body morphology and AP staining after replating

The ESC experiments showed that KDM4 inhibition increased self-renewal-associated phenotypic features; we therefore asked how this response changed during differentiation. At days 10 and 14, DMSO-treated embryoid bodies (EBs) expanded into larger, heterogeneous structures, whereas KDM4i-treated EBs remained smaller and more uniform in both Setdb1 backgrounds (**Fig. 6A-E**). A Detectron2 Mask R-CNN analysis converted this visual phenotype into object classes and showed that KDM4i altered the proportional distribution of asymmetric, cavitated, mixed, and solid morphologies by object number and summed object area (**Fig. 6F**). Extended day-27 imaging and class-resolved analyses showed that these differences persisted (**Supplemental Figs. S6 and S7**).

**Figure 6.**
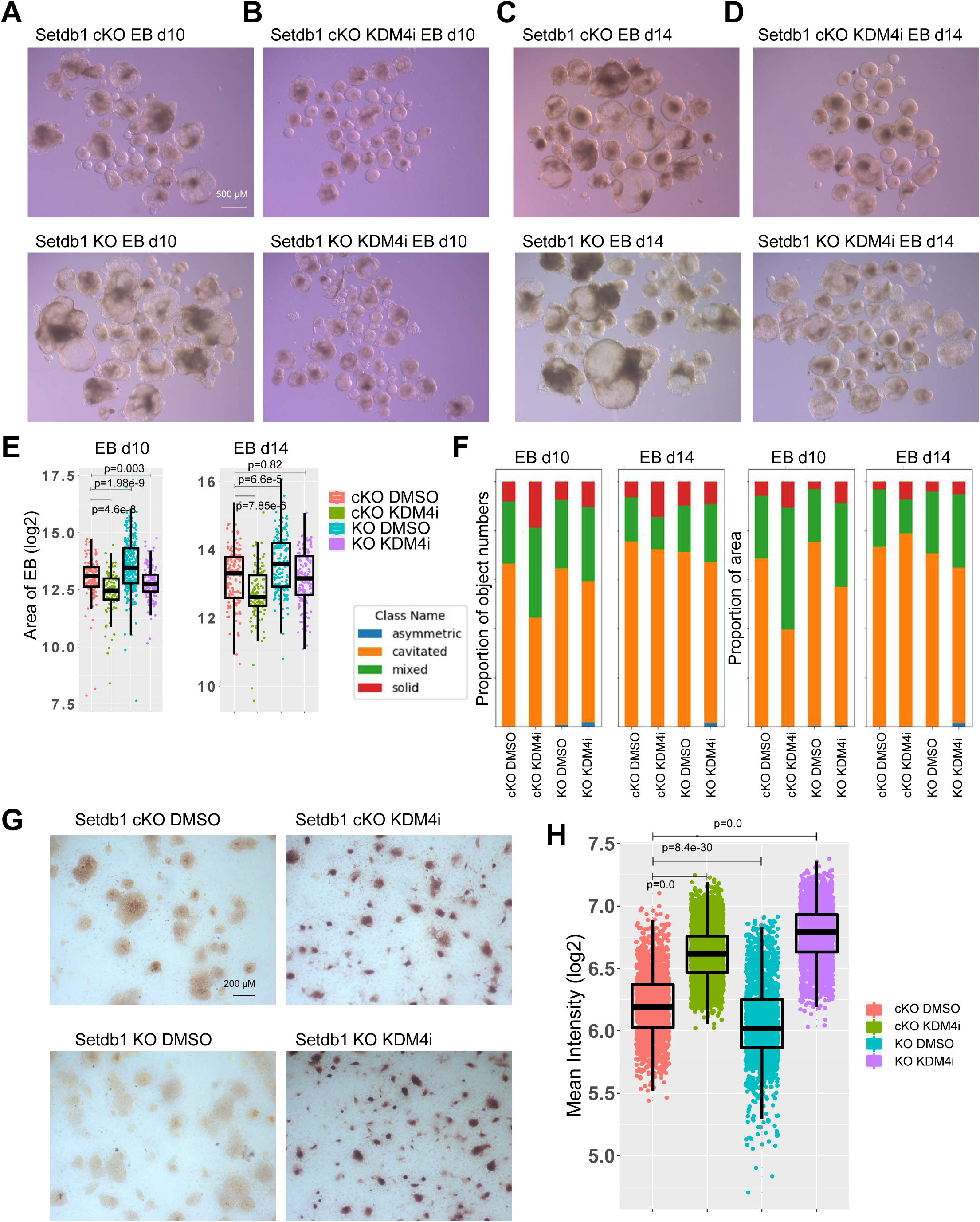
KDM4 inhibition alters embryoid-body morphology and AP staining after replating (A-D) Representative day-10 and day-14 EBs generated from Setdb1 cKO and KO ESCs treated with DMSO or KDM4i. **(E)** Object-level EB area on a log2 scale. **(F)** Detectron2 classification of EB objects by morphology as asymmetric, cavitated, mixed, or solid, shown as proportions of object number and total object area. **(G)** Representative AP staining after prolonged EB differentiation and replating in LIF-containing ESC medium. **(H)** Colony-level mean AP intensity on a log2 scale. Experiments were performed with three to four independent biological replicates. Measurements in E and H represent individual EBs and colonies, respectively, sampled from replicate cultures.

After 27 days of EB differentiation and replating in LIF-containing medium, cells derived from QC6352-treated EBs generated more intensely AP-positive colonies than their DMSO counterparts (**Fig. 6G-H**). This recovery assay measures AP staining after transfer to ESC-supportive culture. The experiments do not distinguish delayed differentiation, reduced proliferation or survival, selection, or altered cell-state composition.

### KDM4 inhibition elicits stage-dependent developmental transcriptional responses that are largely conserved across Setdb1 backgrounds

Day-10 EB RNA-seq revealed that the morphological phenotype was accompanied by a coherent developmental transcriptional state. Three biological replicates per condition clustered by genotype and treatment (**Fig. 7A**). Among 18,853 tested genes, KDM4i induced 793 and repressed 569 genes in cKO EBs and induced 393 and repressed 406 genes in KO EBs (BH-FDR < 0.05 and |log2FC| ≥ 1; **Fig. 7B**). By comparison, Setdb1 loss under DMSO altered 56 induced and 30 repressed genes, whereas the KO-versus-cKO contrast under KDM4i contained 267 induced and 439 repressed genes. The inhibitor therefore remained the dominant perturbation during differentiation, but its response magnitude was reduced in the KO background.

**Figure 7.**
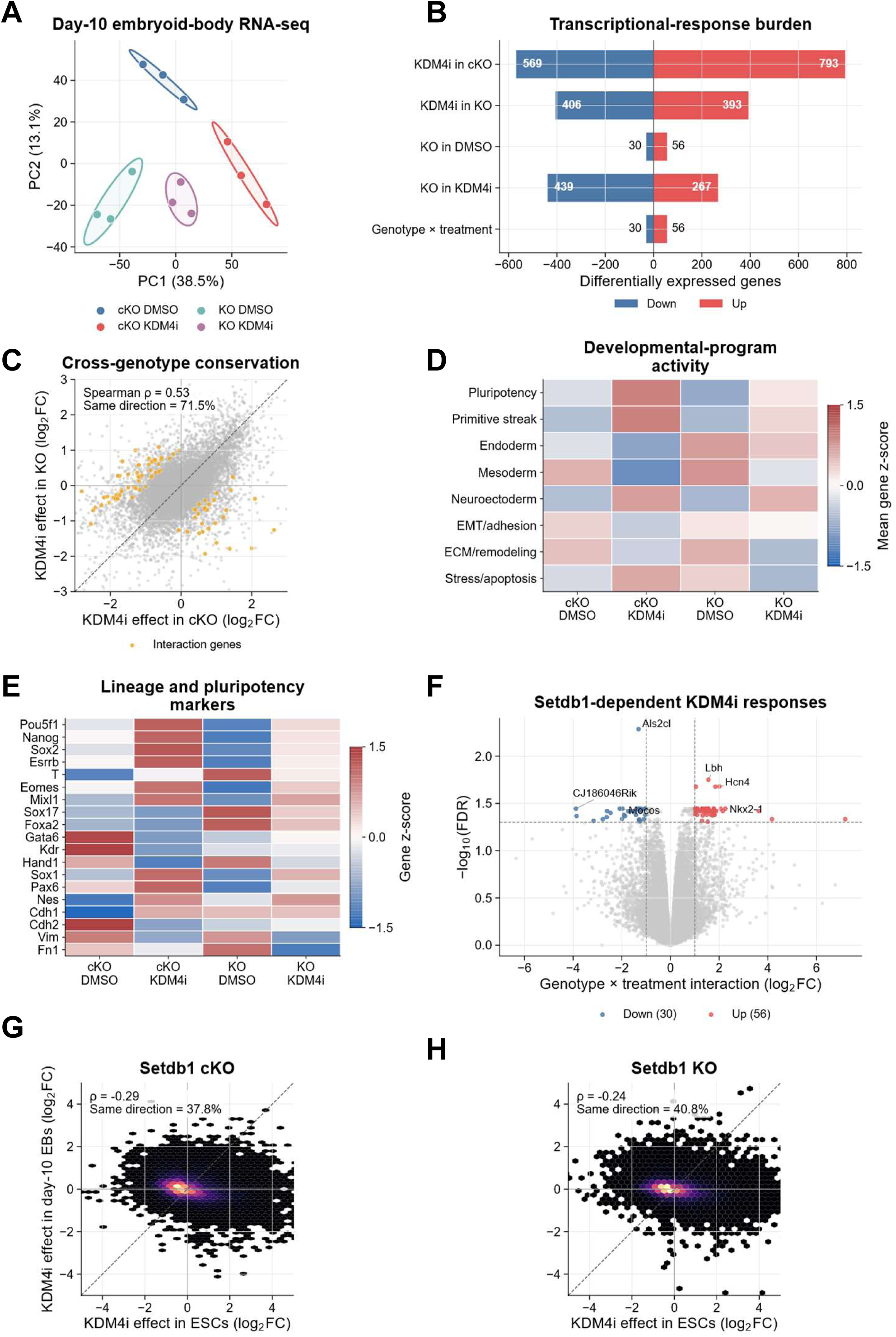
KDM4 inhibition remodels developmental transcription in embryoid bodies. **(A)** PCA of three biological replicates per condition; ellipses are descriptive covariance regions. **(B)** Numbers of upregulated and downregulated genes for five prespecified contrasts: KDM4i versus DMSO in Setdb1 cKO EBs, KDM4i versus DMSO in Setdb1 KO EBs, Setdb1 KO versus cKO under DMSO, Setdb1 KO versus cKO under KDM4i, and the genotype × treatment interaction. Differential expression was defined using edgeR with BH-FDR < 0.05 and |log2FC| ≥ 1. The interaction contrast included 56 positive and 30 negative coefficients. **(C)** Genome-wide KDM4i effects in cKO versus KO EBs; orange points are significant interaction genes. **(D)** Developmental-program scores calculated by averaging replicates within condition, row-standardizing each gene across four condition means, and averaging z-scores within each curated program. **(E)** Gene-wise z-scores for representative pluripotency and lineage markers calculated from the same four condition means. **(F)** Formal interaction volcano plot across all 18,853 tested genes (BH-FDR < 0.05 and |interaction log2FC| ≥ 1; 30 negative, 56 positive). **(G-H)** ESC-versus-day-10-EB KDM4i effects for cKO and KO cells, respectively, across all 17,547 genes common to the independently filtered stage-specific tables without significance preselection. Negative correlations indicate developmental-stage-dependent or reversed response polarity.

KDM4i responses were positively correlated across Setdb1 backgrounds (Spearman ρ = 0.53), with 71.5% of genes responding in the same direction (Fig. 7C). Genotype-by-treatment analysis identified 86 genes with Setdb1-dependent treatment effects, comprising 56 positive and 30 negative interaction coefficients (**Fig. 7B,F**). The interaction bar and volcano highlights are two views of this same restricted set. As in ESCs, the broad response to KDM4 inhibition was therefore largely conserved across Setdb1 states, while formal interaction isolated a small group of genotype-sensitive developmental genes.

Developmental-program and lineage-marker heatmaps placed the conserved KDM4i response in developmental context (**Fig. 7D-E**). KDM4i altered pluripotency, primitive-streak, germ-layer, neuroectodermal, adhesion, extracellular-matrix, and stress-associated programs, indicating broad reorganization of EB state rather than a discrete shift toward a single lineage. Hallmark enrichment across the five factorial contrasts is shown in **Supplemental Fig. S8**.

Comparing KDM4i responses across ESCs and day-10 EBs revealed developmental-stage-dependent transcriptional effects across the 17,547 genes shared by the independently filtered datasets, without significance-based preselection (**Fig. 7G-H**). Responses were modestly anticorrelated in both cKO (ρ = −0.29; 37.8% same direction) and KO cells (ρ = −0.24; 40.8% same direction), consistent with distinct transcriptional responses to KDM4 inhibition in pluripotent and differentiating cells. This stage-dependent pattern was observed despite close clustering of EB replicates and positive concordance of EB responses across Setdb1 genotypes. Together, these findings indicate that the transcriptional consequences of KDM4 inhibition depend on developmental state. In conjunction with the ESC, chromatin, repeat, and morphogenesis results, these findings support a model in which KDM4 inhibition alters selected cellular features in Setdb1-depleted ESCs while producing distinct transcriptional and morphological responses during differentiation.

## DISCUSSION

Our findings argue against viewing SETDB1 and KDM4 enzymes as a simple, globally opposed writer-eraser pair. QC6352 shifted colony organization and AP staining in SETDB1-deficient ESCs toward selected control features without establishing restored self-renewal, yet its transcriptional effect was much larger than the effect of SETDB1 loss itself. The factorial design separates a broad shared QC6352 response from a restricted Setdb1-sensitive component. The M1 and M3 response-reversal modules are candidates for the original biological question, but their contribution to the cellular phenotype is unresolved.

Consistent with prior studies^5, 15, 33, 34^, SETDB1 deficiency impaired ESC colony organization and alkaline phosphatase activity, whereas QC6352 changed these features in depleted and intact cells. These observations motivate further tests of methyltransferase and demethylase contributions to cell state^35, 36^. Across 10,599 KDM4i-responsive genes, 73.2% were significant and directionally concordant in both SETDB1 genotypes and a further 24.9% were significant in one genotype and concordant in the other; only 1.8% were directionally discordant. SETDB1 status modifies the edge of a largely genotype-conserved KDM4i response rather than determining its bulk magnitude.

Independent Kdm4a, Kdm4b, and Kdm4c knockdowns produced correlated but non-equivalent RNA programs, with overlapping responses accompanied by substantial member-specific outputs. These genetic perturbations provide context for the broad pharmacologic response but are not equivalent to simultaneous KDM4-family inhibition. At the chromatin level, KDM4 inhibition reorganized H3K9me3 and H3K36me3 between promoters and gene bodies. This positional redistribution is consistent with the broader principle that domain organization can influence transcriptional competence^37, 38^, but it should not be conflated with an absolute genome-wide gain in a histone mark. The gene-body-to-promoter index measures the log2 gene-body-to-promoter ratio; consequently, an increased ratio can coexist with reduced total H3K36me3 signal.

The repeat analysis similarly refines a uniform heterochromatin-restoration model. SETDB1-dependent H3K9 methylation is a central component of retrotransposon control^5, 39^, and compensatory chromatin mechanisms may help preserve genome stability^13, 40^. Nevertheless, KDM4 inhibition did not simply restore H3K9me3 across all repeat families. Selected ERV families showed reduced H3K9me3 together with increased repeat RNA, whereas the family-wide association between H3K9me3 remodeling and transcription was weak. Importantly, both the chromatin and RNA responses were reproducible across SETDB1 genotypes, indicating that the dominant repeat-family response is also largely conserved across Setdb1 backgrounds. These data associate QC6352 exposure with changes in repeat chromatin^37, 40, 41^, but caution against assuming that demethylase inhibition uniformly reinforces silencing or that H3K9me3 change alone predicts repeat transcription.

The developmental data reveal an additional layer of context dependence. KDM4 inhibition generated smaller, more uniform embryoid bodies, shifted morphological states away from cavitated and mixed structures, and increased recovery of alkaline-phosphatase-positive colonies after replating. These findings are most consistent with altered or delayed morphogenesis and retention of recoverable self-renewal-associated potential, rather than proof that pluripotency is continuously preserved during differentiation. Day-10 embryoid-body RNA-seq reinforced this interpretation: KDM4i responses remained substantially conserved across SETDB1 genotypes, while developmental programs were broadly reorganized rather than redirected toward a single lineage. Moreover, ESC and embryoid-body KDM4i effects were modestly anticorrelated, supporting stage-dependent differences in response polarity.

Together, the data establish broad QC6352-associated transcriptional and relative chromatin-distribution responses, with a restricted subset of transcriptional effects modified by Setdb1 status. Selected cellular features improve within Setdb1-depleted cultures.

## MATERIALS & METHODS

### ESC culture, Setdb1 depletion, and KDM4 inhibition

Mouse ESCs were cultured using established conditions with the modifications described below^42, 43^. Tamoxifen-inducible Setdb1 conditional-knockout (cKO) ESCs^15^ were provided by Matthew Lorincz and Yoichi Shinkai. Setdb1-depleted (KO) ESCs were generated from the conditional line by tamoxifen treatment for four days. Cells were maintained on gelatin-coated dishes in DMEM supplemented with 15% fetal bovine serum, leukemia inhibitory factor (LIF; ESGRO), and 1.5 µM CHIR9901 at 37°C in a humidified CO2-controlled incubator. Cells were routinely washed with PBS, dissociated with trypsin, and passaged before reaching overconfluence. QC6352 was used as the KDM4-family inhibitor (KDM4i); DMSO served as vehicle. ESC experiments comparing genotype and treatment used a 2 × 2 design consisting of Setdb1 cKO DMSO, cKO KDM4i, KO DMSO, and KO KDM4i.

### KDM4-family shRNA knockdown and qRT-PCR

Independent shRNAs targeting Kdm4a, Kdm4b, or Kdm4c were cloned into the pGreenPuro shRNA expression lentivector (System Biosciences); an shRNA targeting firefly luciferase in the same vector, shLuc, served as the negative control. The pGreenPuro vector expresses the shRNA from the H1 promoter and contains copGFP and puromycin-resistance markers for identification and selection of transduced cells. Multiple independent shRNA sequences were evaluated for each Kdm4 paralog to distinguish reproducible target-associated effects from construct-specific responses.

Lentiviral particles were generated in 293T cells by cotransfection of each pGreenPuro-shRNA construct with the psPAX2 packaging plasmid and pLP/VSV-G envelope plasmid using Lipofectamine 2000. Virus-containing medium was collected 24 to 48 h after transfection and used to transduce R1 ESCs. Beginning 24 h after transduction, cells were selected with puromycin at 1 μg/ml. Cells expressing shLuc or the indicated Kdm4-targeting shRNA were collected after selection for qRT-PCR and RNA-seq analysis.

Total RNA was isolated using Qiagen RNeasy kit, and was reverse-transcribed. Quantitative PCR was performed with primers specific for Kdm4a, Kdm4b, Kdm4c, and Actb. Relative transcript abundance was calculated by the 2^−ΔΔCt method using Actb as the reference gene and shLuc as the calibrator. qRT-PCR bars show the mean relative quantity of the included measurements, and error bars denote standard deviation.

### Alkaline-phosphatase imaging and colony quantification

Alkaline-phosphatase (AP)-stained ESC colonies were quantified using ColonyQuant, an automated image-analysis toolkit for colony segmentation, AP-intensity measurement, and morphometric analysis^44^. The ColonyQuant software and documentation are available through the KidderLab repository (https://github.com/KidderLab/ColonyQuant). Segmentation outputs were visually reviewed to confirm appropriate colony detection and background exclusion.

### Embryoid-body differentiation, replating, and image analysis

Embryoid-body differentiation was initiated by transferring cells to low-attachment dishes to promote suspension aggregation and by withdrawing LIF. EBs were imaged at the reported differentiation days. For the recovery assay, differentiated EBs were dissociated and replated in LIF-containing ESC medium before AP staining.

### Detectron2 segmentation and EB morphology classification

EB instances were manually annotated with polygon masks in makeSense.ai and exported in COCO JSON format. A Detectron2 Mask R-CNN instance-segmentation model^45^ was trained on the annotated images and applied through the Detectron2 prediction workflow. Predicted objects were assigned to symmetric, asymmetric, cavitated, mixed, or solid morphology classes. Object masks were used to calculate area and class composition. Model output was visually inspected on held-out images before batch quantification. Class fractions were calculated separately by object number and by summed object area. Because multiple objects can occur in one image or culture, these summaries were interpreted descriptively unless culture-level replication was explicitly modeled.

### RNA-seq library preparation and read quantification

Poly(A)-selected RNA-seq libraries were prepared using the NEBNext Ultra II RNA Library Prep Kit for Illumina and sequenced on an Illumina platform. Sequence reads were aligned to the mouse mm10 reference genome using Bowtie2 with the processing pipeline used for the deposited datasets. Gene-level expression was quantified by assigning mapped reads to RefSeq gene models and calculating raw read counts for statistical analysis. Library-size-normalized RPKM values and genomic coverage tracks were also generated for expression visualization and genome-browser inspection.

### Factorial differential-expression analysis of Setdb1 and KDM4i experiments

ESC and day-10 EB experiments were analyzed separately in edgeR^46^. Lowly expressed genes were removed with filterByExpr using the four genotype-treatment groups or the full design matrix. Library-composition factors were estimated with trimmed mean of M-values normalization (calcNormFactors). A negative-binomial generalized linear model was fitted with genotype, treatment, and genotype × treatment terms, with cKO and DMSO as reference levels. The ESC analysis used robust quasi-likelihood dispersion estimation and glmQLFit(…, robust = TRUE); the EB analysis used the quasi-likelihood workflow without the robust option. Five prespecified contrasts were extracted: KDM4i in cKO, KDM4i in KO, Setdb1 KO versus cKO under DMSO, Setdb1 KO versus cKO under KDM4i, and the genotype × treatment interaction. Benjamini-Hochberg (BH) correction was applied within each contrast.

ESC differential expression required BH-FDR < 0.05 and |log2 fold change| ≥ 0.58. EB differential expression required BH-FDR < 0.05 and |log2 fold change| ≥ 1.

### RNA-seq PCA, reciprocal validation, and interaction modules

PCA was performed on normalized logCPM values after selecting the 500 most variable genes for ESCs and the 5,000 most variable genes for day-10 EBs. Ellipses are descriptive covariance regions and are not confidence intervals. To assess cross-genotype replication of the ESC KDM4i response, genes were ranked by raw P value in the discovery-genotype contrast and evaluated with the independently fitted KDM4i contrast from the other genotype. Fractions with the same effect direction and fractions independently passing BH-FDR < 0.05 and |log2FC| ≥ 0.58 were calculated for the top 50, 100, 200, 500, 1,000, 2,000, 5,000, and 9,000 genes. At each threshold, the random-gene reference was the 2.5^th^-97.5th percentile from 300 size-matched draws without replacement, using separate deterministic seeds for the reciprocal analyses.

The ESC KDM4i-responsive union comprised genes meeting the ESC differential-expression threshold in either genotype. Genes were classified as significant in both genotypes with concordant direction, significant in one genotype with concordant direction in the other, or discordant. Formal interaction genes were defined independently from the genotype × treatment contrast using BH-FDR < 0.05 and |interaction log2FC| ≥ 0.58. For expression modules, logCPM values for the 188 interaction genes were standardized within gene across the ESC libraries. K-means clustering was performed with k = 4, 50 random starts, and seed 20260724. Modules were assigned descriptive labels from their condition-mean profiles. The EB interaction set was defined across all 18,853 tested genes with BH-FDR < 0.05 and |interaction log2FC| ≥ 1.

### Individual KDM4-family knockdown RNA-seq analysis

Each KDM4-targeting shRNA construct was independently profiled by RNA-seq with two biological replicates and contrasted with the shLuc control. Construct-level effects were summarized computationally across the independently analyzed hairpins targeting Kdm4a, Kdm4b, or Kdm4c. The shRNA constructs were not physically pooled before RNA-seq. A core-consensus gene required an absolute mean hairpin log2 fold change ≥ 0.5, complete agreement in effect direction among the target-specific hairpins, and differential-expression support in at least half of those hairpins. Between-hairpin standard deviation was retained as a measure of construct heterogeneity. Aggregate target-level differentially expressed genes were defined using BH-FDR < 0.05 and |log2FC| ≥ 0.5. Shared and member-specific sets were defined from exact intersections of the three significant-gene sets. Genome-wide similarity was quantified from pairwise correlations of aggregate target-level log2 fold changes.

### Gene-set enrichment and inferred pathway activity

GO and Hallmark gene-set enrichment analyses used ranked gene statistics and BH correction across the tested gene sets. For interaction-module GO Biological Process analysis, gene symbols were mapped to Entrez identifiers and clusterProfiler enrichGO was applied separately to each module using all expressed ESC genes with an Entrez mapping as the universe, BH adjustment, gene-set sizes of 10-500 genes, and no significance prefilter before reporting. EB Hallmark enrichment used the signed square root of the edgeR quasi-likelihood F statistic to preserve effect direction while incorporating statistical strength.

PROGENy activities were inferred with decoupleR^47^ using the mouse PROGENy network and the multivariate linear-model method. Input genes were ranked by the signed statistic −log10(raw P value) × log2FC. BH correction was performed within each contrast across the 14 pathways. Heatmaps use one symmetric activity scale; markers denote BH-FDR < 0.05. Pathways were clustered for display, whereas perturbations remained in the prespecified biological order. Correlations between aggregate knockdown and KDM4i activity profiles were calculated across the 14 pathway scores. These transcriptome-derived activities are inferential and do not constitute direct biochemical measurements.

### ChIP-seq library preparation and primary processing

ChIP-seq experiments followed established procedures with modifications^48, 49^. Antibodies were H3K4me3 (Millipore 17-614), H3K27ac (Abcam ab4729), H3K9me3 (Abcam ab8898), H3K36me3 (Abcam ab9050), and H4K20me3 (Abcam ab9053). ESCs were crosslinked with 1% formaldehyde for 10 min, and chromatin from approximately four million cells was immunoprecipitated with 2 µg antibody. Libraries were generated with NEBNext Ultra II end-repair/dA-tailing reagents, Illumina adapters, and Phusion high-fidelity PCR, and were sequenced on an Illumina platform. Reads were aligned to mm10 with Bowtie2^50^, converted to BED6, deduplicated, and summarized as library-size-normalized reads per base per million mapped reads (RPBM). Two biological replicates were analyzed per condition and histone mark.

Enriched regions were called relative to matched input DNA with SICER using a 200-bp window, 400-bp gap, and FDR 0.001; SICER-compare used FDR < 0.001 and fold change > 1.5 where pairwise domain comparisons were required. Counts were interpreted within mark because focal and broad-domain region calls are not directly comparable. Genomic-feature composition was calculated from peak annotations: gene body combined exon and intron, distal/intergenic corresponded to intergenic sequence, and Other combined 3′UTR, 5′UTR, transcription termination site, miRNA, ncRNA, pseudogene, rRNA, and snoRNA annotations. Replicate-level signal PCA used normalized genomic signal for each mark. Genome-wide reproducibility was evaluated by Spearman correlation of normalized coverage in fixed genomic bins.

### Expression-stratified metagene profiles and gene-body-to-promoter-index analysis

Genes were divided into quartiles using ESC expression. Metagene profiles extended from 3 kb upstream of the transcription start site (TSS), across a scaled gene body from TSS to transcription end site (TES), to 3 kb downstream. KDM4i/DMSO or KO/cKO ratios were calculated within biological replicate before averaging. Solid lines represent replicate means and ribbons span replicate values. For display, profile means and ribbon boundaries were smoothed after ratio calculation with a centered nine-bin rolling mean. **Figure 4** profiles used input-normalized ChIP signal. The **Supplemental Figure S4** sensitivity analysis used library-normalized ChIP signal without division by input DNA.

Promoter and gene-body measurements were obtained from the PausingIndex pipeline, in which promoter signal is stored separately from gene-body signal. Finite positive transcript-level measurements were retained, transcript isoforms were collapsed to the median per gene and replicate, and biological replicates were averaged within condition. Gene-body-to-promoter index was defined as log2(gene-body signal/promoter signal); KDM4i-induced Δ gene-body-to-promoter index was the KDM4i value minus matched DMSO within genotype. Analyses included genes with complete measurements across the required conditions. Phase portraits plotted promoter log2(KDM4i/DMSO) against gene-body log2(KDM4i/DMSO). Cross-genotype concordance used Spearman correlation and the fraction of genes with the same Δ-index direction. Paired Wilcoxon tests compare matched gene-level responses and are descriptive rather than biological-replicate inference. The ratio is a descriptive positional index, not a measurement of domain expansion.

### Repeat-family H3K9me3 and RNA integration

Repeat annotations were derived from the mm10 RepeatMasker catalogue. Repeat copies were intersected with H3K9me3 signal and with repeat-overlapping RNA measurements using the bed2refseq, RPKM, density-calculation, and merge workflows. The integrated analysis was restricted to 49 repeat families represented in both biological replicates and all four genotype-treatment conditions. For each repeat copy, density denotes H3K9me3 ChIP-seq density and RPKM/count values denote repeat-overlapping RNA signal. Family-level H3K9me3 effects were calculated as log2 ratios of condition-specific median copy values; repeat RNA was summarized as a family-aggregate count response. Biological-replicate concordance was evaluated from replicate-specific family effects. H3K9me3-RNA coupling and cross-genotype conservation used Spearman correlation and directional agreement. Repeat copies are not independent biological replicates, and multi-mapping and within-family heterogeneity limit locus-level interpretation.

### Developmental-program scores and ESC-EB response comparisons

For EB developmental-program heatmaps, normalized logCPM values were averaged across the three biological replicates within each genotype-treatment condition. Each gene was then standardized across the four condition means, and curated program scores were calculated as the arithmetic mean of the constituent gene z-scores. The lineage-marker heatmap displays the individual standardized gene values. For ESC-EB comparisons, KDM4i log2 fold changes were matched by gene across the independently filtered stage-specific contrast tables. All common genes were included without differential-expression preselection. Spearman correlation and same-direction fraction were calculated separately for cKO and KO cells.

## DECLARATIONS

### ETHICS APPROVAL AND CONSENT TO PARTICIPATE

Not applicable

### CONSENT FOR PUBLICATION

All authors have read and approved the final version of this manuscript.

## Data Availability

The sequencing data from this study have been submitted to the NCBI Gene Expression Omnibus (GEO). Accession information will be provided upon public release.

## Biological replication, statistical tests, and reporting

Biological triplicate RNA-seq datasets were generated for each genotype-treatment condition in ESCs and day-10 EBs. Biological duplicate ChIP-seq datasets were generated for H3K9me3, H3K36me3, H3K4me3, H3K27ac, and H4K20me3 in Setdb1 cKO and KO ESCs treated with DMSO or KDM4i. Unless stated otherwise, multiplicity was controlled with the Benjamini-Hochberg method and all tests were two-sided. Exact analysis populations, thresholds, sample sizes, and denominators are stated in the figure legends and Methods. Gene-level, repeat-copy-level, colony-level, and EB-object-level distributions do not increase the number of independent biological replicates; tests over these units are identified as descriptive. Custom analysis scripts used fixed random seeds for stochastic procedures, and analysis outputs retain unrounded effect estimates, raw P values, and adjusted P values.

## COMPETING INTERESTS

The authors declare no conflict of interest.

## AUTHORS’ CONTRIBUTIONS

B.L.K. conceived of the study, designed and assisted with the experiments, analyzed the sequencing data, and drafted the manuscript. R.G. carried out experiments and assisted with revising the manuscript.

## ACKNOWLEDGEMENTS

We thank Zengquan Yang for kindly providing the KDM4 inhibitor (QC6352), obtained from MCE, and for helpful discussions. This work utilized the Wayne State University High Performance Computing Grid for computational resources.

## FUNDING

Wayne State University; Barbara Ann Karmanos Cancer Institute [P30 CA022453— Cancer Center Support Grant].

## Notes

### Competing Interest Statement

The authors have declared no competing interest.

