## Supplemental information for "KDM4 inhibition alters H3K9me3 distribution and developmental responses in SETDB1-deficient embryonic stem cells"

Running title: KDM4 inhibition and H3K9me3 in SETDB1-deficient ESCs.

[*]Correspondence:

Benjamin L. Kidder

**KEYWORDS**

H3K9me3; Setdb1; KDM4; embryonic stem cells; pluripotency; lineage specification; ERV silencing; epigenetic regulation; transcriptional dynamics; RNA-Seq; ChIP-Seq

### Supplemental Figures

#### Figure S1. Dose-dependent ESC morphology and alkaline-phosphatase responses to KDM4 inhibition

(A) Representative bright-field images of Setdb1 cKO and KO ESCs treated with the indicated KDM4i concentrations. (B) Representative bright-field images of R1 ESCs treated with DMSO or 100–1,000 nM KDM4i. (C) Representative alkaline-phosphatase (AP)-stained R1 ESC cultures treated with DMSO or the indicated KDM4i concentrations. (D) Colony-level mean AP intensity on a log2 scale across the KDM4i dose series. Boxes denote the interquartile range and center lines denote medians. Scale bars are shown in the panels. (E) Uncropped immunoblot images corresponding to the cropped blots shown in Figure 1C-D. ESC morphology and AP assays were performed with three to four independent biological replicates. Colony measurements were sampled from replicate cultures.

#### Figure S2. Protein validation and transcriptome-derived pathway responses following KDM4-family knockdown

(A) Immunoblot validation of KDM4A, KDM4B, and KDM4C protein depletion following the indicated shRNAs. HSC70 serves as the loading control. Representative blots are shown. (B) Transcriptome-derived PROGENy activity estimates for Kdm4a, Kdm4b, and Kdm4c knockdown responses summarized across independently analyzed shRNAs. Rows show the 14 PROGENy pathways and columns show the aggregate responses for Kdm4a, Kdm4b, and Kdm4c knockdown. Color denotes the signed decoupleR activity statistic on a symmetric scale; asterisks denote BH-FDR < 0.05. (C) Representative GO gene-set enrichment results for genes induced or repressed in the aggregate Kdm4a, Kdm4b, or

Kdm4c knockdown responses derived from independently analyzed shRNAs. Color denotes normalized enrichment score, with red indicating positive enrichment and blue indicating negative enrichment. Point size denotes the indicated significance category. (D) Uncropped immunoblot images corresponding to the cropped blots shown in (A).

**Figure S3. ChIP-seq enrichment and reproducibility metrics**

Quality and reproducibility summaries for H3K27ac, H3K4me3, H3K36me3, H3K9me3, and H4K20me3 ChIP-seq. (A) Numbers of enriched regions identified in each biological replicate and condition, shown in thousands. Counts are interpreted within each histone mark because focal and broad-domain calls are not directly comparable. (B) Genomic-feature composition of annotated enriched regions. Promoter denotes promoter-associated regions, gene body combines exon and intron, distal/intergenic denotes intergenic sequence, and Other combines 3′UTR, 5′UTR, transcription termination site, miRNA, ncRNA, pseudogene, rRNA, and snoRNA annotations. (C) PCA of replicate-level normalized genomic signal for each histone mark. Colors denote genotype-treatment conditions and symbols denote biological replicates. (D) Pairwise Spearman correlations of normalized coverage in fixed genomic bins.

**Figure S4. ChIP-only positional profiles and gene-body-to-promoter-index distributions for H3K36me3 and H4K20me3**

Positional analyses calculated from library-normalized ChIP RPBM signal without division by input DNA. (A) Expression-stratified H3K36me3 metagene fold-change profiles for KDM4i versus DMSO in Setdb1 cKO and KO cells and for Setdb1 KO versus cKO under DMSO. Genes were divided into ESC-expression quartiles; profiles extend from 3 kb upstream of the transcription start site through the scaled gene body to 3 kb downstream of the transcription end site. Lines show replicate means and ribbons span replicate values. (B) ECDFs of the H3K36me3 gene-body-to-promoter index, defined as log2(gene-body signal/promoter signal), for the same comparisons. Annotations report paired gene counts and median comparison-minus-reference shifts. (C-D) Corresponding H4K20me3 metagene profiles and gene-body-to-promoter-index ECDFs.

**Figure S5. Additional repeat-class H3K9me3 profiles**

Element-centered H3K9me3 heatmaps for MERVL-int and MT2_Mm repeat copies across Setdb1 cKO and KO ESCs treated with DMSO or KDM4i. Rows represent eligible repeat copies grouped by response pattern, and columns show signal across the repeat-centered interval for each genotype-treatment condition. These profiles extend the repeat-family examples in Figure 5; mapping and eligible-copy criteria are described in Methods.

**Figure S6. Extended embryoid-body morphology after Setdb1 loss and KDM4 inhibition**

Day-27 embryoid-body morphology across the four genotype-treatment conditions. (A-B) Representative bright-field images of Setdb1 cKO and KO EBs treated with DMSO or KDM4i. (C) Individual EB area on a log2 scale. Boxes denote the interquartile range and center lines denote medians. (D-E) Representative Detectron2 instance-segmentation outputs for DMSO- and KDM4i-treated EBs, respectively. (F) Proportional distribution of Detectron2-classified EB morphologies, calculated by EB number and summed EB area, for asymmetric, cavitated, mixed, and solid classes. The scale bar is shown in panel A. EB experiments were performed with three to four independent biological replicates. Individual EBs were sampled from replicate cultures.

**Figure S7. Day-27 embryoid-body segmentation and class-resolved principal-component summaries**

(A) Detectron2-segmented day-27 EB images from Setdb1 cKO DMSO, cKO KDM4i, KO DMSO, and KO KDM4i cultures. Colored masks identify individual predicted objects and labels indicate assigned morphology classes. (B) PCA of class-resolved EB measurements at days 10, 14, and 27, calculated separately from summed EB area and EB number. Colors denote morphology class and plotting symbols denote genotype-treatment group.

**Figure S8. Hallmark pathway responses in day-10 embryoid bodies**

Signed Hallmark gene-set enrichment across KDM4i effects in Setdb1 cKO and KO EBs, Setdb1 KO effects under DMSO and KDM4i, and the formal genotype-by-treatment interaction. Rows show the 20 pathways with the strongest combined evidence across contrasts. Color denotes normalized enrichment score on a symmetric scale, with red indicating positive enrichment and blue indicating negative enrichment; asterisks denote BH-FDR < 0.05. Complete normalized enrichment scores, raw P values, BH-FDR values, gene-set sizes, and leading-edge genes are provided in Supplemental Table S4.

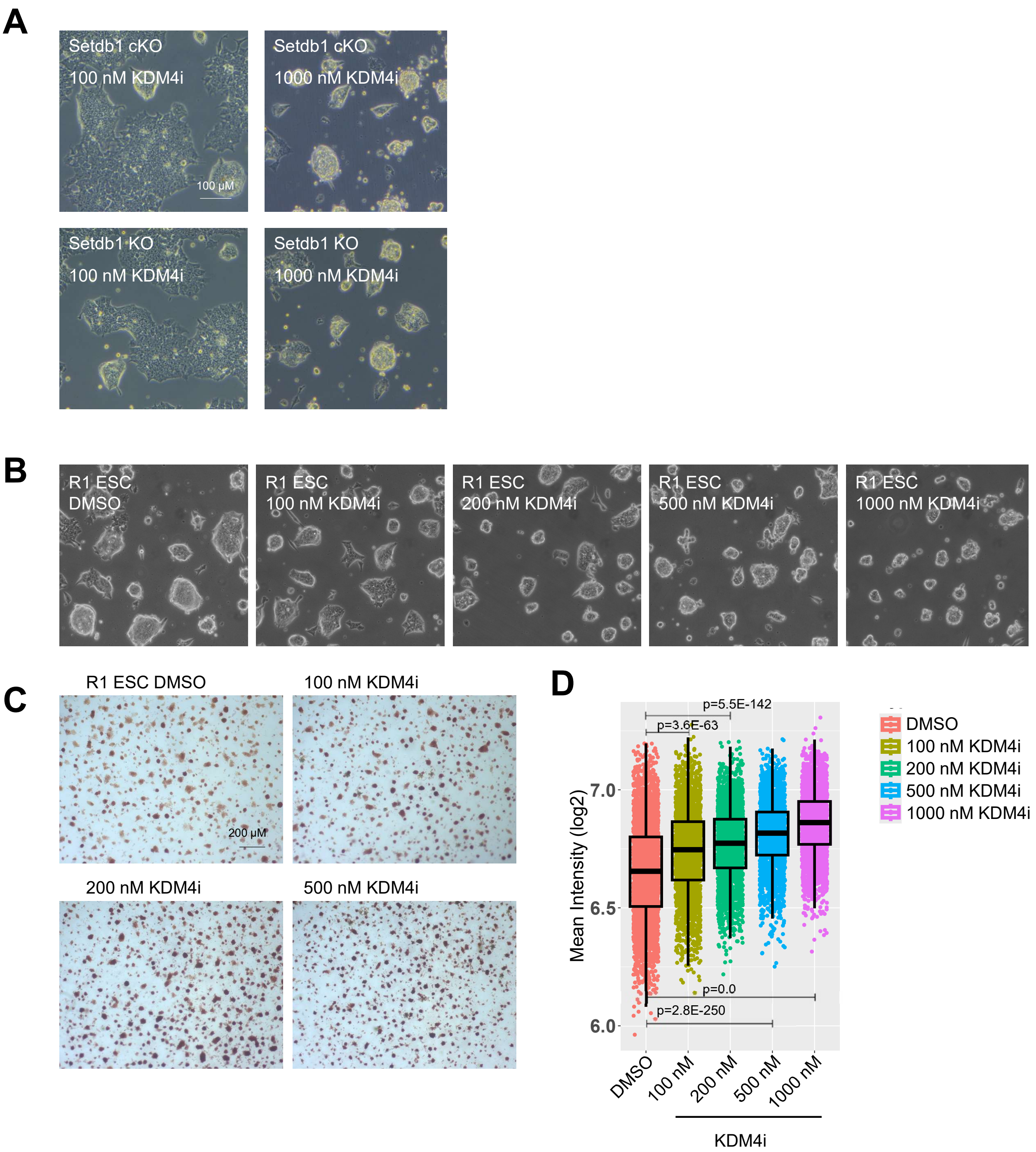
A
Setdb1 cKO
100 nM KDM4i
100 μM
Setdb1 cKO
1000 nM KDM4i
Setdb1 KO
100 nM KDM4i
Setdb1 KO
1000 nM KDM4i
B
R1 ESC
DMSO
R1 ESC
100 nM KDM4i
R1 ESC
200 nM KDM4i
R1 ESC
500 nM KDM4i
R1 ESC
1000 nM KDM4i
C
R1 ESC DMSO
100 nM KDM4i
200 μM
200 nM KDM4i
500 nM KDM4i
D
p=5.5E-142
p=3.6E-63
p=0.0
p=2.8E-250
Mean Intensity (log2)
7.0
6.5
6.0
DMSO
100 nM
200 nM
500 nM
1000 nM
KDM4i
DMSO
100 nM KDM4i
200 nM KDM4i
500 nM KDM4i
1000 nM KDM4i

Figure S1

E

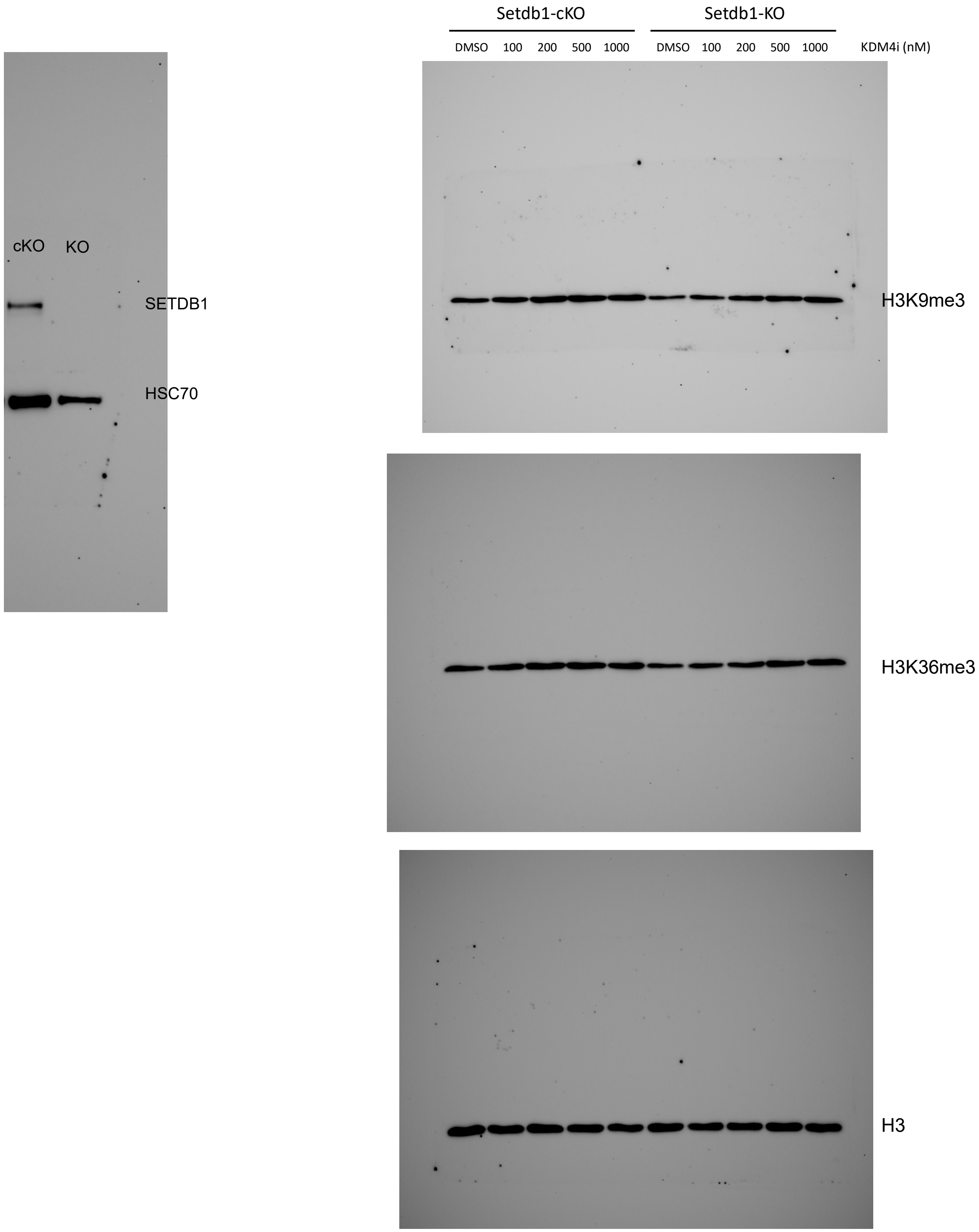
cKO
KO
SETDB1
HSC70
Setdb1-cKO
Setdb1-KO
DMSO
100
200
500
1000
DMSO
100
200
500
1000
KDM4i (nM)
H3K9me3
H3K36me3
H3

Figure S1

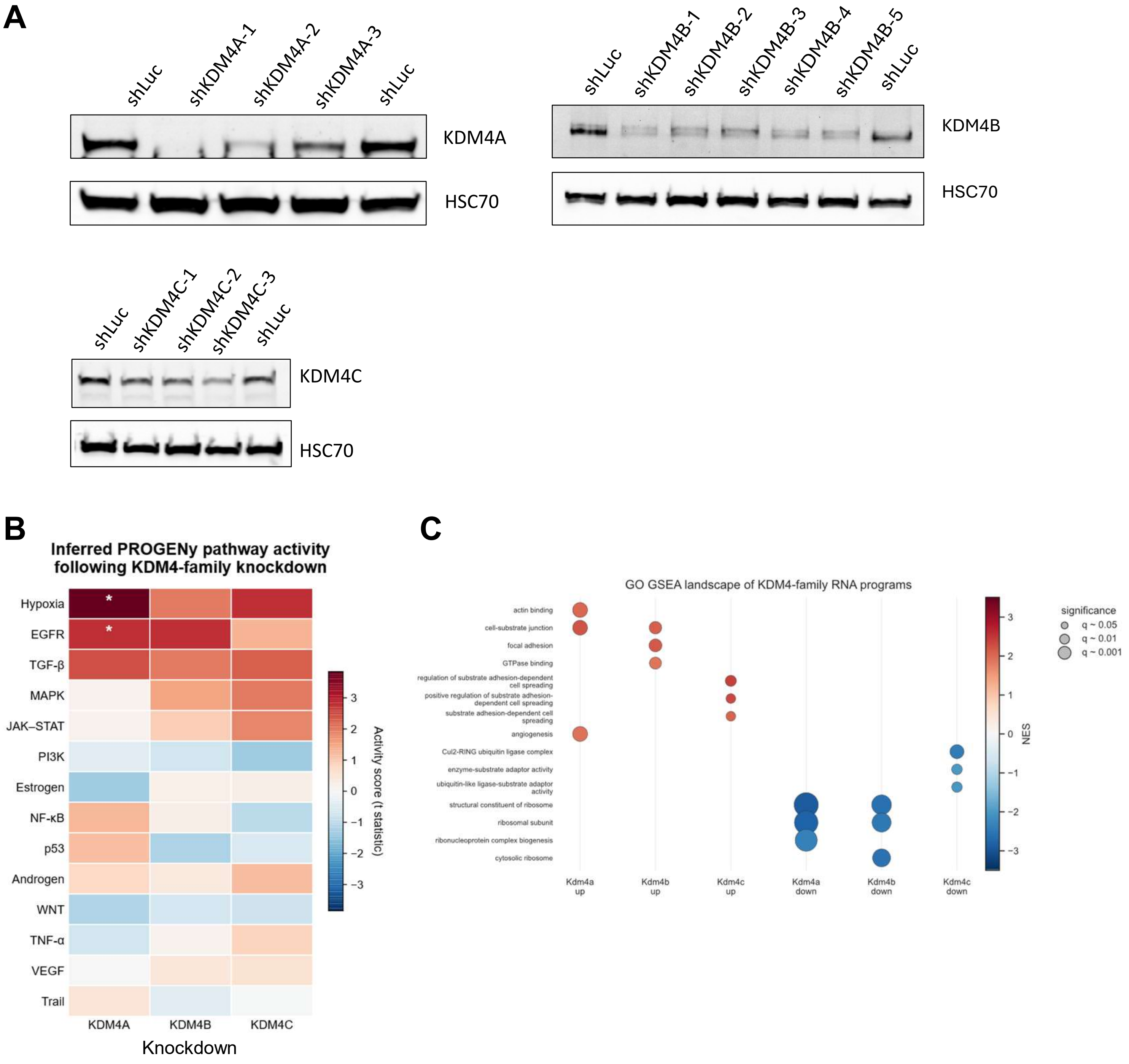
A
shLuc
shKDM4A-1
shKDM4A-2
shKDM4A-3
shLuc
KDM4A
HSC70
shLuc
shKDM4B-1
shKDM4B-2
shKDM4B-3
shKDM4B-4
shKDM4B-5
shLuc
KDM4B
HSC70
shLuc
shKDM4C-1
shKDM4C-2
shKDM4C-3
shLuc
KDM4C
HSC70
B
Inferred PROGENy pathway activity
following KDM4-family knockdown
Hypoxia
EGFR
TGF-β
MAPK
JAK–STAT
PI3K
Estrogen
NF-κB
p53
Androgen
WNT
TNF-α
VEGF
Trail
KDM4A
KDM4B
KDM4C
Knockdown
Activity score (t statistic)
C
GO GSEA landscape of KDM4-family RNA programs
actin binding
cell-substrate junction
focal adhesion
GTPase binding
regulation of substrate adhesion-dependent cell spreading
positive regulation of substrate adhesion-dependent cell spreading
substrate adhesion-dependent cell spreading
angiogenesis
Cul2-RING ubiquitin ligase complex
enzyme-substrate adaptor activity
ubiquitin-like ligase-substrate adaptor activity
structural constituent of ribosome
ribosomal subunit
ribonucleoprotein complex biogenesis
cytosolic ribosome
Kdm4a up
Kdm4b up
Kdm4c up
Kdm4a down
Kdm4b down
Kdm4c down
NES
significance
q ~ 0.05
q ~ 0.01
q ~ 0.001

**Figure S2**

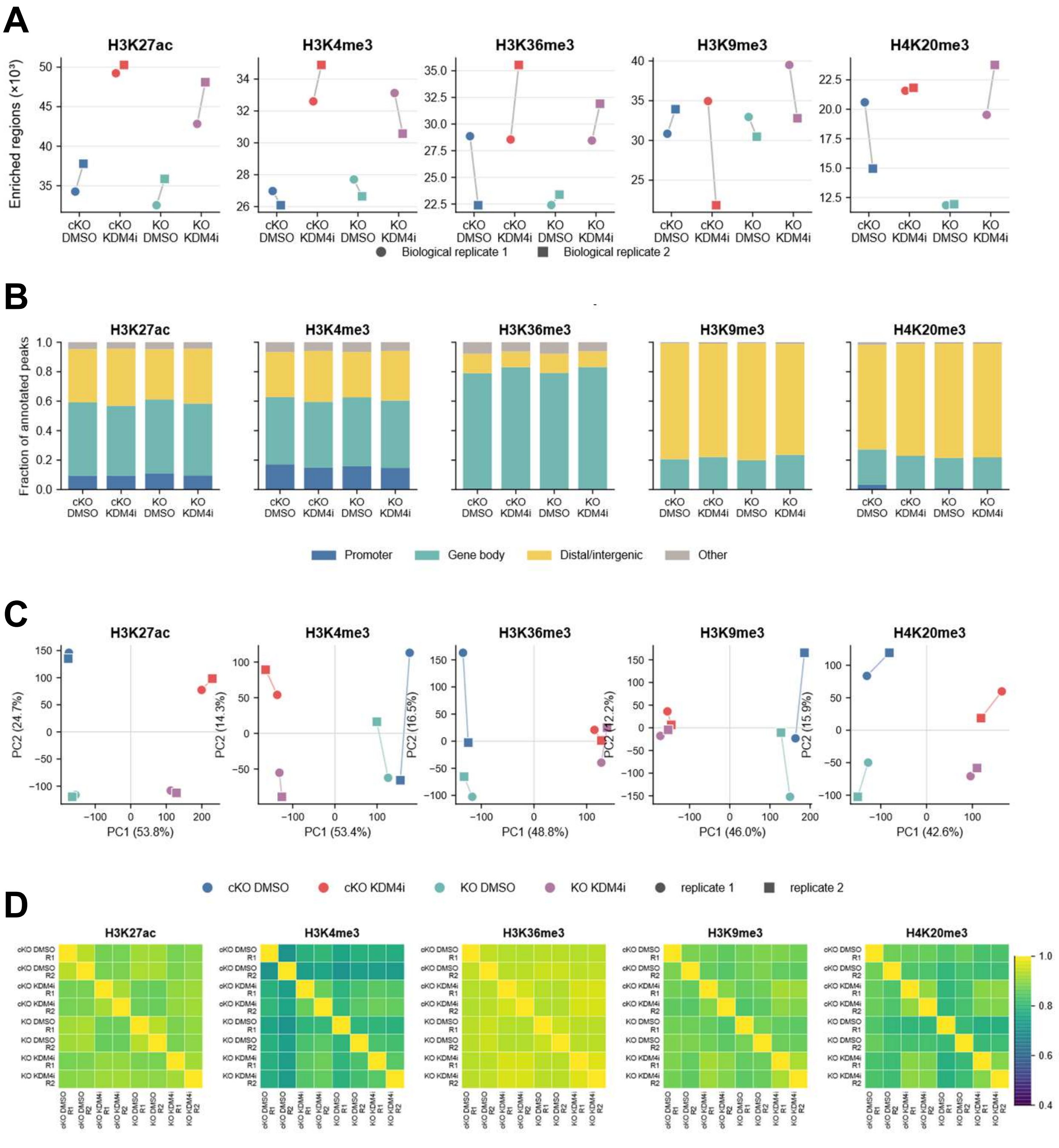
A
H3K27ac
H3K4me3
H3K36me3
H3K9me3
H4K20me3
Enriched regions (×10³)
cKO DMSO
cKO KDM4i
KO DMSO
KO KDM4i
Biological replicate 1
Biological replicate 2
B
Fraction of annotated peaks
Promoter
Gene body
Distal/intergenic
Other
C
PC1 (53.8%)
PC2 (24.7%)
PC1 (53.4%)
PC2 (14.3%)
PC1 (48.8%)
PC2 (16.5%)
PC1 (46.0%)
PC2 (12.2%)
PC1 (42.6%)
PC2 (15.9%)
cKO DMSO
cKO KDM4i
KO DMSO
KO KDM4i
replicate 1
replicate 2
D
Spearman ρ

Figure S3

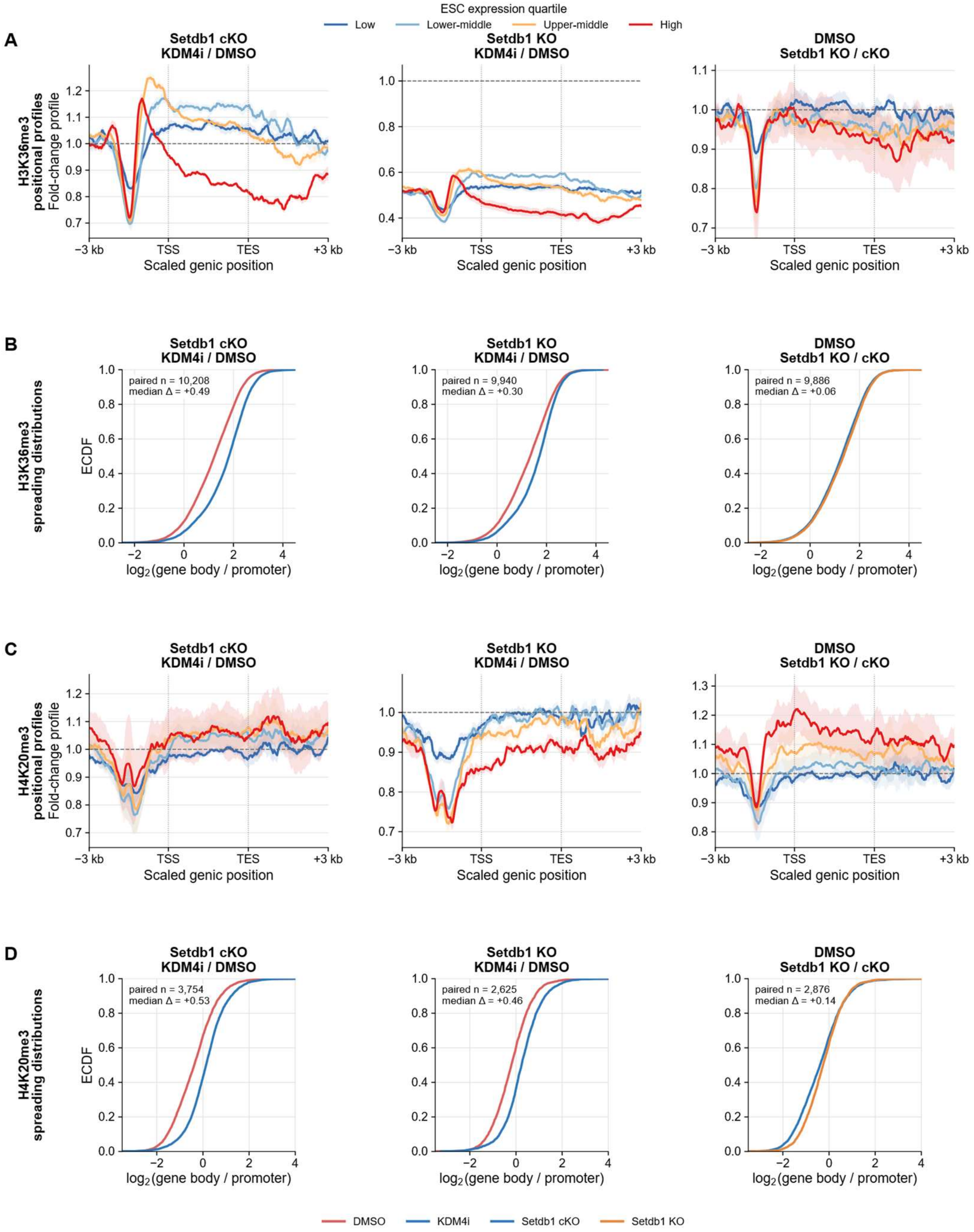
ESC expression quartile
Low
Lower-middle
Upper-middle
High
A
Setdb1 cKO
KDM4i / DMSO
Setdb1 KO
KDM4i / DMSO
DMSO
Setdb1 KO / cKO
H3K36me3
positional profiles
Fold-change profile
Scaled genic position
−3 kb
TSS
TES
+3 kb
B
H3K36me3
spreading distributions
ECDF
paired n = 10,208
median Δ = +0.49
paired n = 9,940
median Δ = +0.30
paired n = 9,886
median Δ = +0.06
log2(gene body / promoter)
C
H4K20me3
positional profiles
Fold-change profile
D
H4K20me3
spreading distributions
paired n = 3,754
median Δ = +0.53
paired n = 2,625
median Δ = +0.46
paired n = 2,876
median Δ = +0.14
DMSO
KDM4i
Setdb1 cKO
Setdb1 KO

**Figure S4**

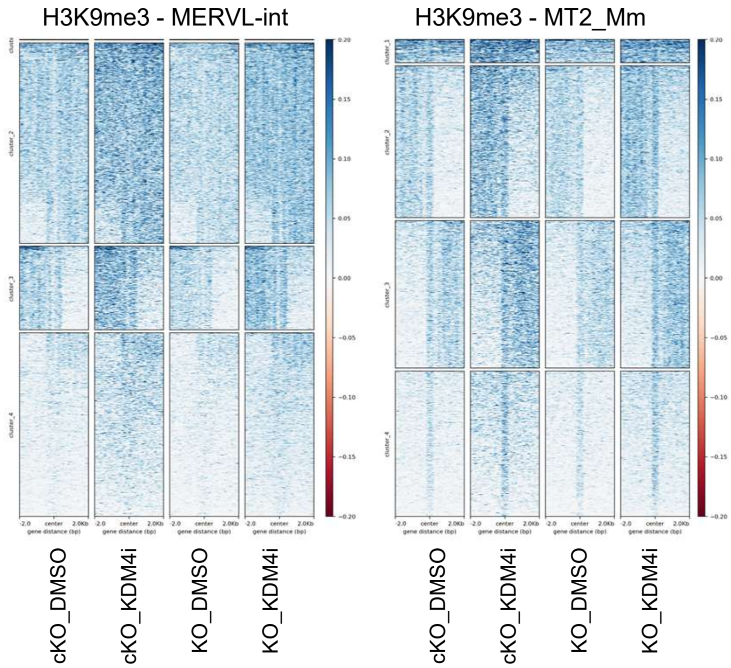

H3K9me3 - MERVL-int
H3K9me3 - MT2_Mm
cKO_DMSO
cKO_KDM4i
KO_DMSO
KO_KDM4i


Figure S5

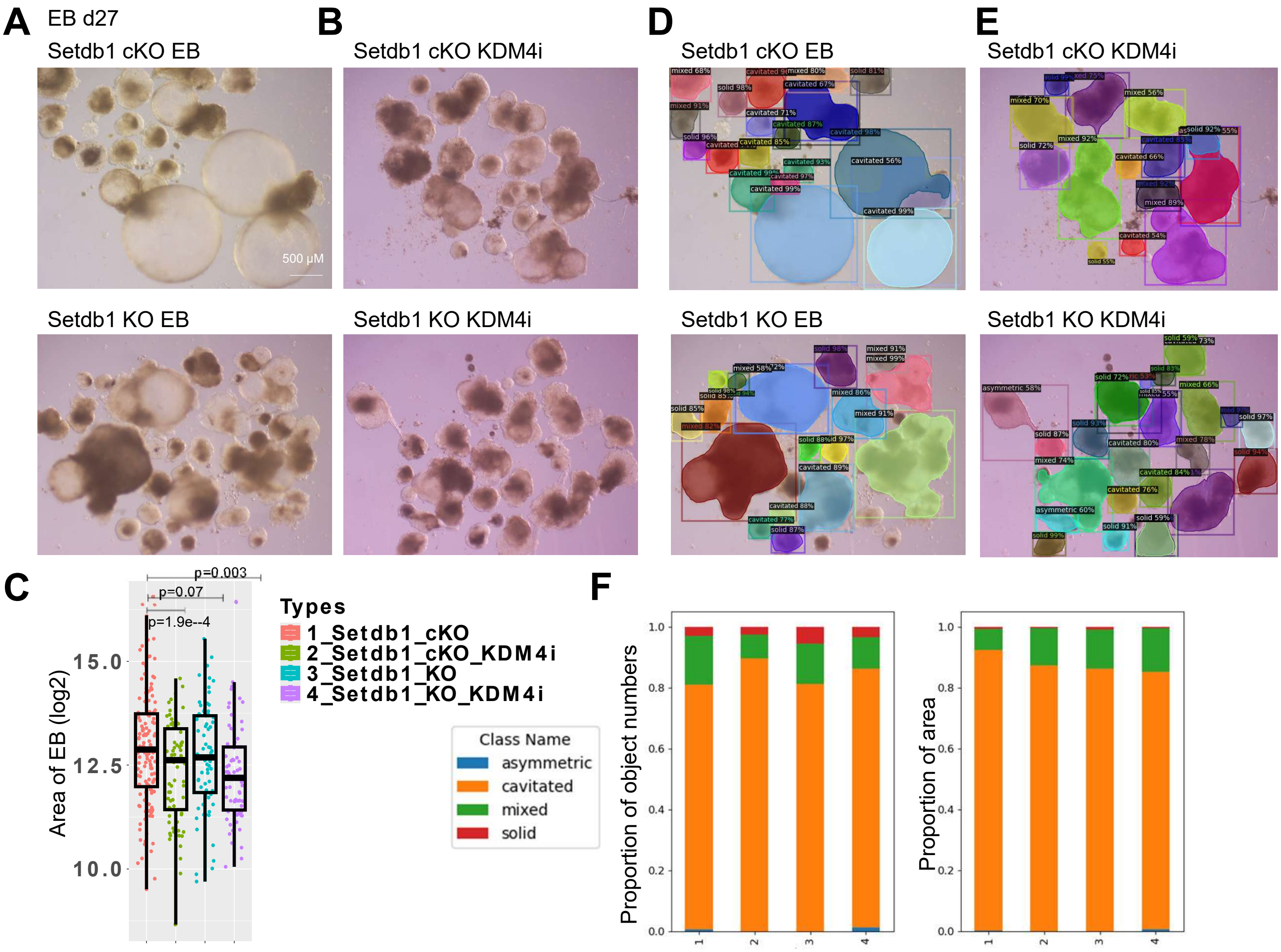
A
EB d27
Setdb1 cKO EB
500 μM
Setdb1 KO EB
B
Setdb1 cKO KDM4i
Setdb1 KO KDM4i
D
Setdb1 cKO EB
Setdb1 KO EB
E
Setdb1 cKO KDM4i
Setdb1 KO KDM4i
C
p=0.003
p=0.07
p=1.9e--4
Area of EB (log2)
15.0
12.5
10.0
Types
1_Setdb1_cKO
2_Setdb1_cKO_KDM4i
3_Setdb1_KO
4_Setdb1_KO_KDM4i
Class Name
asymmetric
cavitated
mixed
solid
F
Proportion of object numbers
Proportion of area

Figure S6

**A**

EB d27

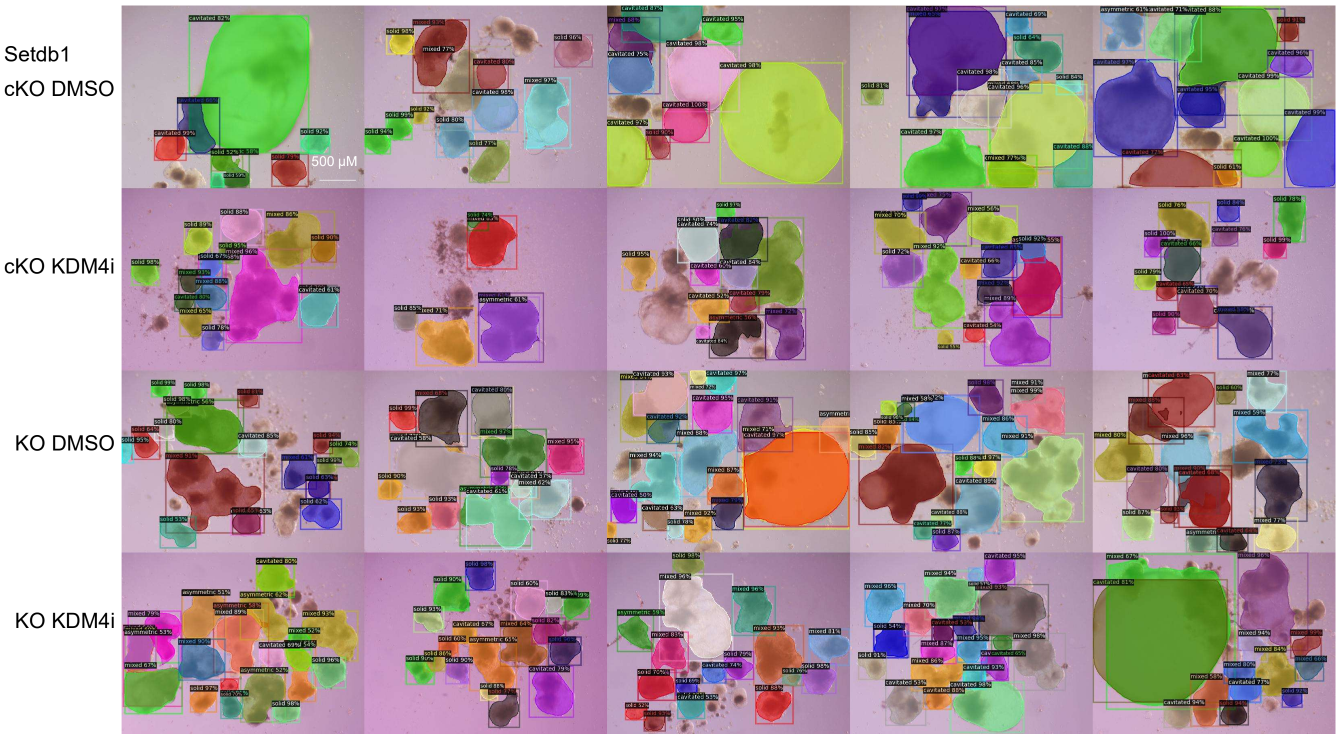


**B**

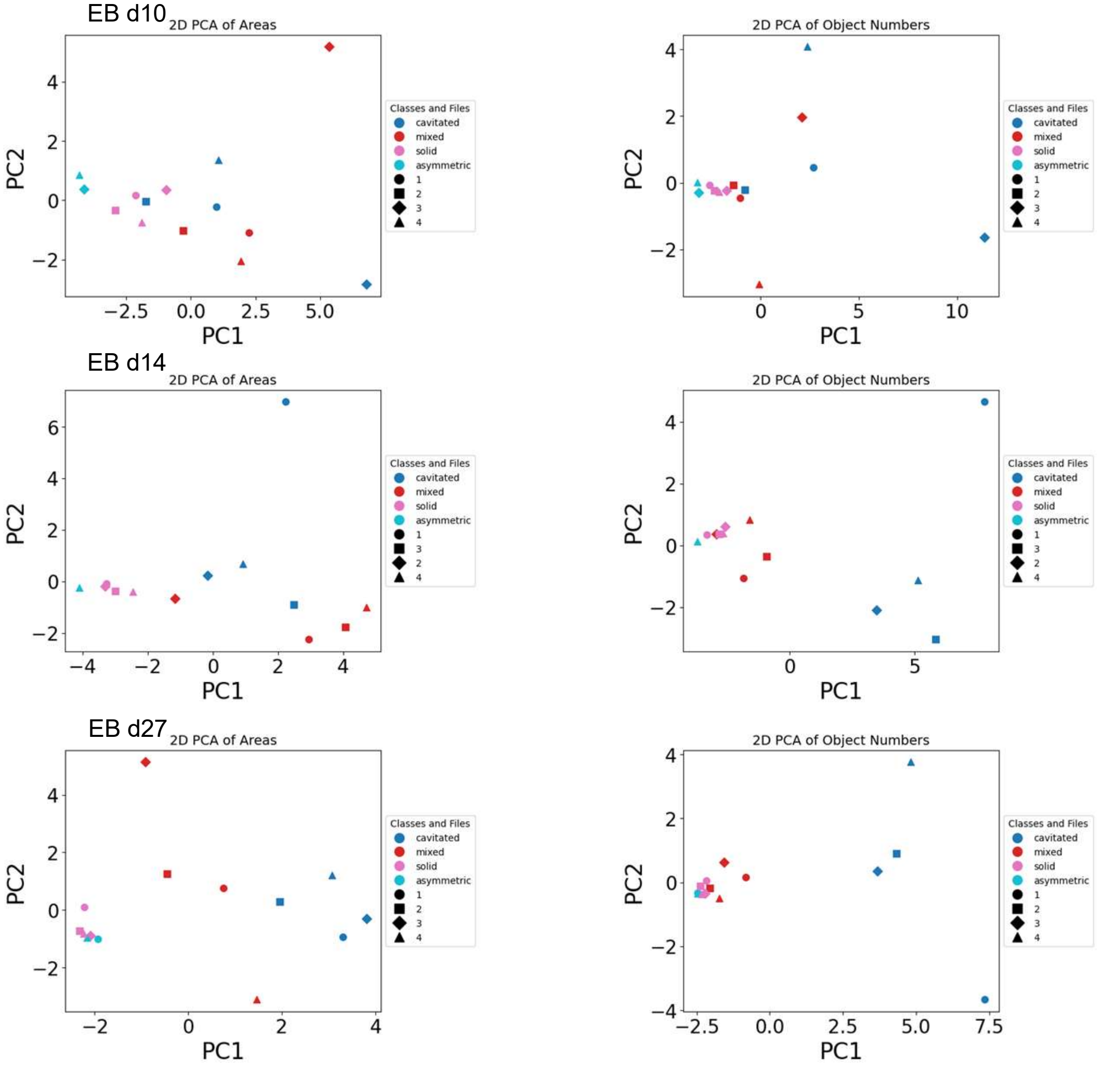


**Figure S7**

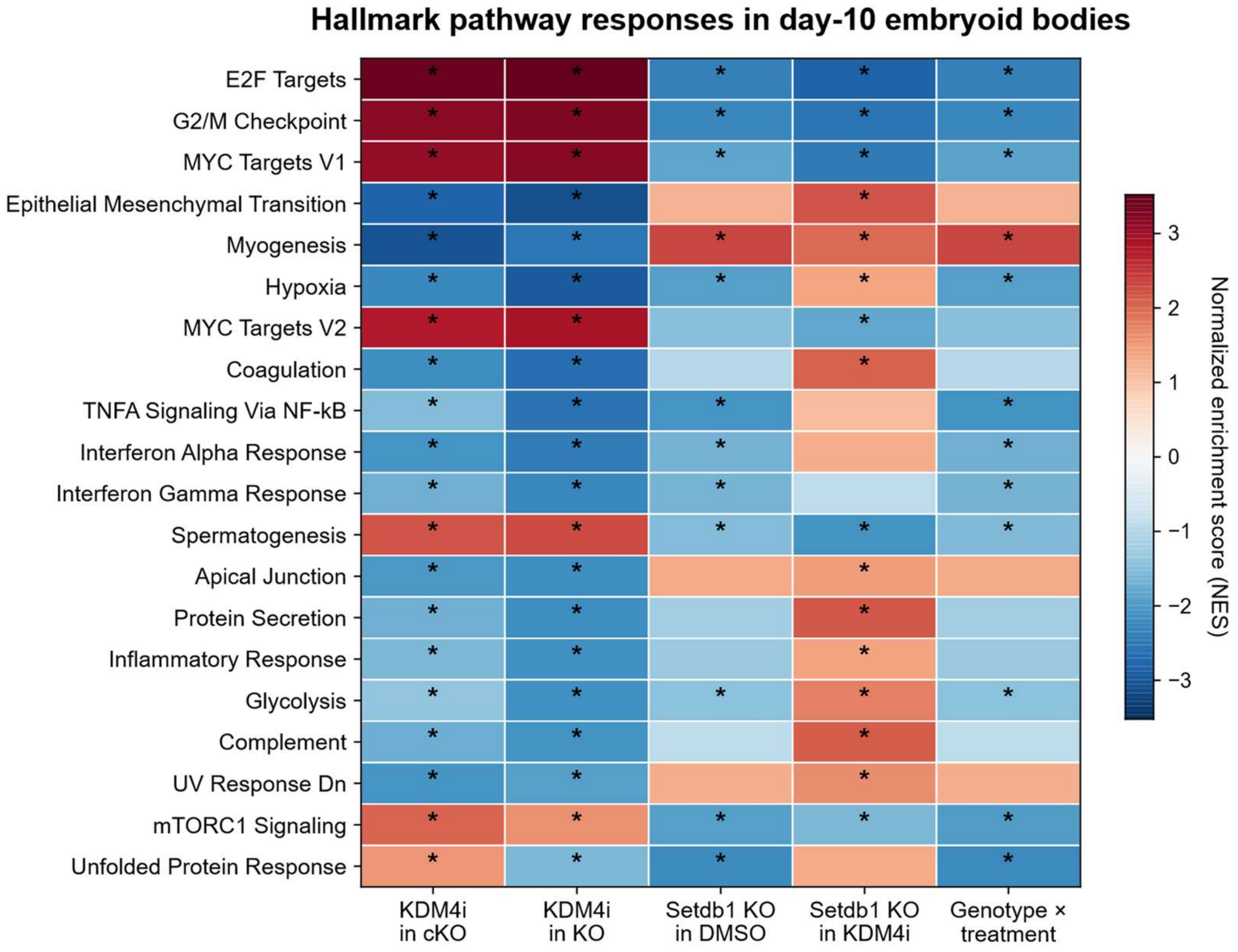
Hallmark pathway responses in day-10 embryoid bodies
E2F Targets
G2/M Checkpoint
MYC Targets V1
Epithelial Mesenchymal Transition
Myogenesis
Hypoxia
MYC Targets V2
Coagulation
TNFA Signaling Via NF-kB
Interferon Alpha Response
Interferon Gamma Response
Spermatogenesis
Apical Junction
Protein Secretion
Inflammatory Response
Glycolysis
Complement
UV Response Dn
mTORC1 Signaling
Unfolded Protein Response
KDM4i in cKO
KDM4i in KO
Setdb1 KO in DMSO
Setdb1 KO in KDM4i
Genotype × treatment
Normalized enrichment score (NES)
3
2
1
0
−1
−2
−3

Figure S8
